# Integrity of the minor spliceosome in the developing mouse hypothalamus determines neuronal subtype composition regulating energy balance

**DOI:** 10.1101/2022.10.04.510883

**Authors:** Alisa K. White, Kyle D. Drake, Alexandra E. Porczak, Gabriela Tirado-Mansilla, Madisen F. Lee, Katery C. Hyatt, Chrissy Chow, Tava DeQuattro, Laura E. Mickelsen, Natale R. Sciolino, Alexander C. Jackson, Rahul N. Kanadia

**Affiliations:** Physiology and Neurobiology Department, University of Connecticut, Storrs, CT 06269, USA; Institute of Systems Genomics, University of Connecticut, Storrs, CT 06269, USA

## Abstract

While gene regulatory networks underlying hypothalamic development are being characterized, minor intron splicing remains unexplored. Here, we used *Nkx2*.*1-Cre* to ablate *Rnu11*, encoding the minor spliceosome-specific U11 snRNA, in the progenitors of the ventral diencephalon (VD), to study minor intron splicing in hypothalamic development and control of energy balance in mice. Loss of U11 resulted in aberrant minor intron splicing, mitotic stalling, apoptosis, and altered neurogenesis. Mutant mice exhibited gross dysgenesis of hypothalamic architecture, while single-cell RNA sequencing (scRNAseq) revealed aberrant composition of neuronal subtypes implicated in feeding and energy balance. Mutant weanlings failed to thrive, followed by rapid weight gain, resulting in obesity. Assessment of energy imbalance and pair-feeding demonstrated that hyperphagia in adult mutants initiates weight gain, and is compounded by metabolic dysfunction, ultimately resulting in obesity. Our findings suggest a key role of minor intron splicing in the developmental patterning of hypothalamic neuronal subtypes underlying energy balance.

## Introduction

Gene expression regulation, through transcription factors and signaling molecules, during hypothalamic development is under active investigation^1^. However, the role of gene regulation through splicing, which occurs co-transcriptionally, remains understudied. The discovery of neurodevelopmental diseases such as Microcephalic Osteodysplastic Primordial Dwarfism Type 1 (MOPD1), Early Onset Cerebellar Ataxia (EOCA), and Inherited Growth Hormone Deficiency (IGHD), that are linked to mutations in components of the minor spliceosome, underscore the importance of minor intron splicing in neural development^2–5^. Here, we explored the role of minor intron splicing in hypothalamic development by employing *Nkx2*.*1-Cre* which has previously been used to perturb hypothalamic development^6,7^. *Nkx2*.*1* is expressed as early as embryonic day (E)10.5 in the ventral diencephalon (VD) progenitors that will give rise to hypothalamic regions controlling feeding (e.g. arcuate (ARC), ventromedial hypothalamus (VMH), dorsomedial hypothalamus (DMH), lateral hypothalamus (LH)), as well as the mammillary nucleus (MM)^8–11^.

The inherent significance of understanding hypothalamic development is that the hypothalamus is essential for the postnatal regulation of physiological and behavioral homeostasis, including arousal, circadian rhythms, energy metabolism, and feeding^12–17^. In fact, several *Nkx2*.*1*-derived hypothalamic regions have been widely studied for their roles in regulating energy balance, including the ARC, VMH, DMH, and LH^15,16,18–21^. Widespread efforts through scRNAseq have provided a rich map of the various neuronal subtypes in the adult mouse hypothalamus^22–29^. However, the developmental programs underlying the production of these subtypes have only recently gained attention^30,31,29,32^. Given that the programs underpinning hypothalamic neuronal subtype identity is established during embryonic development^24,29,31–34^, disruptions in hypothalamic development could alter hypothalamic structure, function, and behavior in postnatal life^30,35–37^. Here, we show that the integrity of the minor spliceosome is essential for hypothalamic development and function.

The mouse genome consists of major introns (>99.5%) and minor introns (<0.5%), that are spliced by their correspondingly named spliceosomes^38^. Although, minor intron-containing genes (666 genes in *mus musculus*^39^) also consist major introns, their expression is uniquely dependent on the minor spliceosome. The minor spliceosome consists of 5 small nuclear RNAs (snRNAs), U11, U12, U4atac, U6atac, and U5, along with over 100 core proteins^38^. To study the unexplored role of minor intron splicing in mouse hypothalamic development, we crossed our conditional knockout (cKO) allele of *Rnu11*^40^, encoding the minor spliceosome specific U11 snRNA, with an *Nkx2*.*1-Cre* driver^9^.

Here, we show that loss of U11 in *Nkx2*.*1*+ progenitor cells, resulted in elevated minor intron retention, leading to impaired cell cycle progression and apoptosis. The developing mutant hypothalamus showed elevated neurogenesis, with concomitant progenitor cell depletion. These developmental insults resulted in mutant mice failing to thrive preweaning (e.g. reduced bodyweight, short stature), followed by postnatal hyperphagia with an extended circadian pattern that resulted in weight gain and obesity. Through scRNAseq from postnatal day (P)30 mice, we observed stark reductions in ARC neuronal subtypes, especially those regulating feeding (*Npy*^+^, *Pomc*^+^), connecting to the hyperphagic behaviors of the obese mutant mice. Obesity was further compounded by reduced energy expenditure and physical activity of the mutant mice. Pair-feeding experiments, where the daily caloric intake of mutants was restricted to match control mice, prevented obesity progression. This demonstrated that hyperphagia is the primary factor responsible for weight gain of mutants. The overall phenotype observed in this mouse model phenocopies key physiological and behavioral sequalae observed in Prader-Willi Syndrome (PWS), independent of the causative genetic insult^41^. Taken together, our findings demonstrate the importance of hypothalamic development in disorders of feeding and reveal the role of minor intron splicing in hypothalamic neuronal subtypes and the control of energy balance.

## Results

### Minor spliceosome is essential from *Nkx2*.*1+* progenitor cell cycle and survival

To inhibit minor intron splicing in the developing hypothalamus, we first crossed a *Rnu11* conditional knockout out (cKO)^40^ mouse to a *Nkx2*.*1-Cre* line^9^. To label the progenitors and their lineage targeted by *Nkx2*.*1-Cre*, we bred in a tdTomato fluorescent reporter allele^42^, to generate our control *(Rnu11*^WT/FLX^:: R26^LSL-tdTomato^;*Nkx2*.*1*-*Cre*^+^) and mutant *(Rnu11*^FLX/FLX^::R26^LSL-tdTomato^;*Nkx2*.*1*-*Cre*^+^) mice. Fluorescent *in situ* hybridization (FISH) for U11, coupled with immunofluorescence (IF) to detect expression of tdTomato in the *Nkx2*.*1-Cre*+ region, was performed at E10.5 through E12.5 to assess U11 loss in the VD (**Fig. 1A-C**). At E10.5, we found successful ablation of U11 in the VD neuroepithelium (**Fig. 1A**). At E11.5, the loss of U11 in the tdTomato+ domain was also observed, except in a subset of cells in the mutant VD, where we observed mosaic expression of U11 despite tdTomato signal (**Fig. 1B**). Similar loss of U11 was observed in the tdTomato^+^ region in the VD of E12.5 embryo, with the lack of U11 loss in the ventral most part of the VD (**Fig. 1C**).

**Figure 1.**
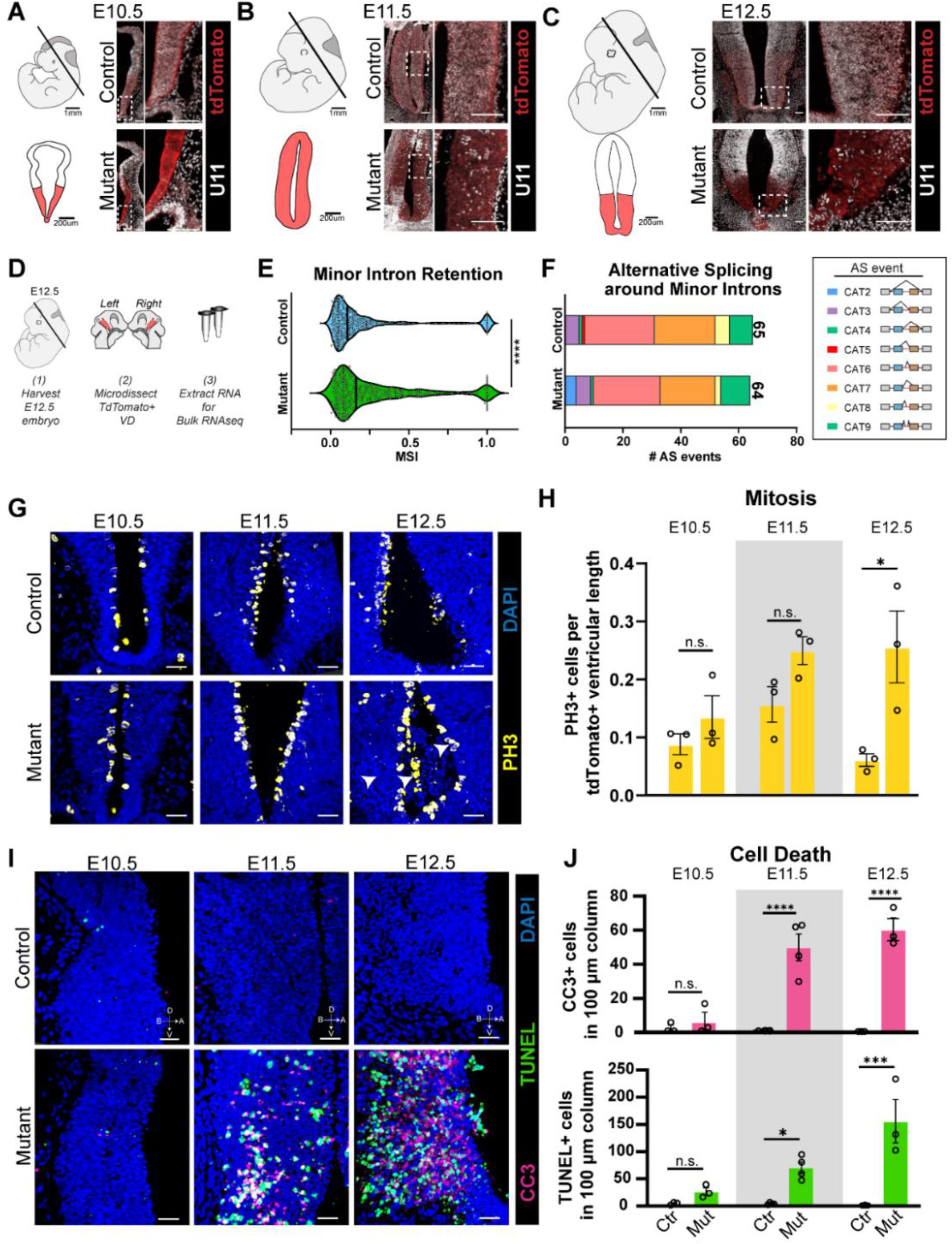
Ablation of *Rnu11* in the ventral diencephalon results in mis-splicing of minor intron-containing genes, cell cycle defects, and cell death. (**a-c**) Fluorescent *in situ* hybridization (FISH) for U11 (white), and immunofluorescence (IF) for RFP to mark the tdTomato reporter (red) on coronal sections of E10.5, E11.5, E12.5 control *(Rnu11*^WT/FLX^:: R26^LSL-tdTomato^;*Nkx2*.*1*-*Cre*^+^) and mutant *(Rnu11*^FLX/FLX^::R26^LSL-tdTomato^;*Nkx2*.*1*-*Cre*^+^) embryos. Scale bars 100µm. (**d**) Schematic of E12.5 ventral diencephalon harvest for bulk RNAseq. (**e**) Violinplot of the median mis-splicing index (MSI) for all (n=409) minor introns that show retention in either control or mutant. (**f**) Stacked bar chart of the detected alternative splicing (AS) events around minor introns in the control (top) and mutant (bottom). (**g**) IF for PH3 (yellow) within the Nkx2.1^+^ region at E10.5, E11.5, E12.5 with DAPI (blue) marking nuclei. Scale bar 30µm. (**h**) Quantification of the number of PH3^+^ mitotic cells over the quantified region (RFP^+^ ventricular length in µm) at E10.5 (n=3), E11.5 (n=4), E12.5 (n=3). (**i**) IF for CC3 (magenta) and TUNEL assay (green) at E10.5, E11.5, and E12.5 with DAPI (blue) marking nuclei. Scale bar 30 µm. (**j**) Quantification of cell death in a 100µm column in the RFP+ region at E10.5 (n=3), E11.5 (n=4), and E12.5 (n=3). Data (**e**) significance was determined by Mann-Whitney U test. ****=*P*<0.0001. Data (**h, j**) represented as mean, with error bars ±SEM. Significance was determined by two-tailed student’s t-tests. n.s.= not significant, *=*P*<0.05, **=*P*<0.01, ***=*P*<0.001, ****=*P*<0.0001 (See also **Supplementary Data Table 9**).

To assess minor intron splicing defects, we next performed bulk RNA-seq on microdissected E12.5 tdTomato+ VD for control (n=3) and mutant (n=3) embryos (**Fig. 1D**)^40,43,44^. The expression of *Nkx2*.*1, Six3*, and *Rax* combined with the absence of *Emx1* and *Foxg1*, confirmed the accuracy of our dissection^45–47^ (**Supplementary Data Table 1**). RNAseq analysis showed significantly elevated minor intron retention in the mutant VD compared to control (**Fig. 1E**). Amongst the 74 MIGs with significantly increased minor intron retention, we found several MIGs involved in ‘cell cycle’ (including *Dctn3, Mau2, Cctn1, Pten*) and ‘neuron projection development’ (including *Spg11, Myh10, Slc9a6, Vash2*). Moreover, AS that skips the upstream and downstream exons flanking the minor intron (CAT2), was elevated in the mutant VD relative to controls (**Fig. 1F, blue**).

Given that MIGs regulating cell cycle showed elevated minor intron retention, we next investigated whether mutants had alterations in mitotic proliferation. At E10.5 and E11.5, similar number of PH3+ VD in mitosis was observed in the mutant and controls (**Fig. 1G, H**). An increase PH3+ VD cells of the mutant was observed E12.5 (**Fig. 1G, H**), suggesting a possible increase in progenitor cells in the mutant developing hypothalamus. Upon further inspection, we observed multinucleated cells that appeared to fail to complete mitosis, including cells in the ventricular space with pyknotic nuclei (**Fig. 1G, white arrowheads**). We therefore assessed whether these cells might be undergoing apoptosis by performing IF for cleaved caspase 3 (CC3), coupled with terminal deoxynucleotidyl transferase dUTP nick end labeling (TUNEL). Indeed, we observed significantly elevated CC3+ and TUNEL+ cells at E11.5, which continued to E12.5 in the mutant VD relative to the control (**Figure 1I, J**). Altogether we show the primary defect of U11 ablation is elevated minor intron retention, defective cell cycle, and apoptosis in the developing VD.

### Ablation of U11 in the VD reduced progenitor subtypes and elevated neurogenesis

The elevated apoptosis in the mutant VD led us to explore the distribution of various progenitor subtypes in the mutant VD. As the VD has a complex three-dimensional (3D) structure, we utilized whole mount *in situ* hybridization (WISH) on hemisected control and mutant embryos at E12.5 to visualize the entire VD. In mutants, we observed a decrease in *Nkx2*.*1* and an absence of U11 in the *Nkx2*.*1*+ VD (**Fig. 2A**). A similar, yet condensed pattern of expression of *Shh*, a known morphogen regulating VD patterning was observed in mutant embryos (**Fig. 2A**). Further a reduction in the expression domain of *Lhx6*, was found at the intrahypothalamic diagonal, forming a weak boundary of pre-hypothalamic and pre-thalamic structures (**Fig. 2A**). Next, we interrogated markers of subregions within the VD corresponding to progenitor pools that give rise to distinct structures of the postnatal hypothalamus. *Irx5*, know to largely give rise to the supramammillary nucleus (SUM)^10^, had a similar expression pattern within the retromammillary, albeit slightly reduced, in the mutants (**Fig. 2A**). In contrast, *Foxb1*, marking the mammillary nucleus, appeared greatly condensed in its expression pattern in mutant embryos (**Fig. 2A**). Finally, *Lef1*, marking the premammillary nucleus (PM), and *Pomc*, marking the ARC, revealed reductions in their staining patterns in the VD (**Fig. 2A**), demonstrating that ablation of *Rnu11* disrupted the spatial pattern of progenitor subtypes in the developing VD.

**Figure 2.**
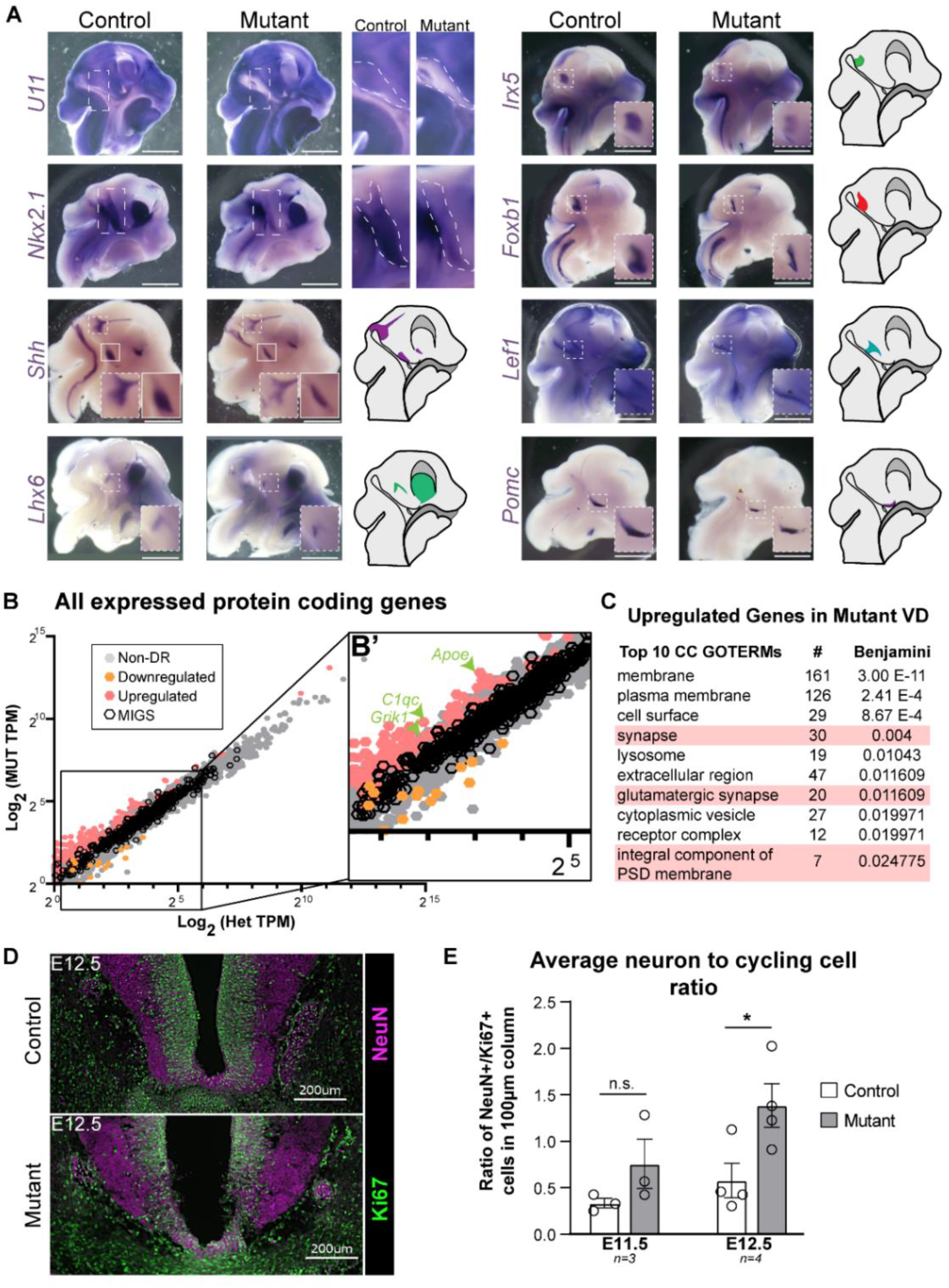
Impaired minor intron-splicing results in pre-hypothalamic progenitor cell depletion, and increased neuron production. (**a**) Whole-mount *in situ* hybridization of E12.5 control and mutant hemisected embryos for U11 and Nkx2.1, patterning marker (*Shh*), and progenitor subdomain markers (*Lhx6, Irx5, Foxb1, Lef1*, and *Pomc*). Scale bar 1mm. Zoomed insets highlighting ventral diencephalon expression patterns. (**b**) Scatterplot of all expressed (1TPM in either genotype) protein coding genes. (**b’**) Zoomed inset of scatterplot with green arrow heads pointing to some upregulated genes in the mutant ventral diencephalon enriching for the GOTerm ‘synapse’. (**c**) Top 10 cellular component GOTerms significantly enriched for by the upregulated genes in the mutant ventral diencephalon. Those enriching for neuron-related terms highlighted in pink. (**d**) Immunofluorescence for Ki67 (green), NeuN (magenta) in control and mutant coronal ventral diencephalon at E12.5. (**e**) Quantification of the ratio of neurons (NeuN+) to cycling progenitor cells (Ki67+) in a 100µm column at E11.5 (n=3) and E12.5 (n=4). Scale bar 200µm. Data (**e**) represented as mean, with error bars ±SEM. Significance was determined by two-tailed student’s t-tests. n.s.= not significant, *=*P*<0.05 (See also **Supplementary Data Table 9**).

This disruption in the number of progenitor subtypes in the mutant VD, led us to investigate the role of *Rnu11* ablation on neurogenesis. Global gene expression analysis between control and mutant showed 331 upregulated genes in the mutant VD (**Fig. 2B,B’**), with enrichment for neuronal-related GOTerms, including “synapse” (30 genes), “glutamatergic synapse” (20 genes), and “integral component of PSD membrane” (7 genes) (**Fig. 2C**). Based on this enrichment, we hypothesized that the mutant VD is undergoing elevated neurogenesis which was investigated by IF for Ki67 (cycling cells), coupled with NeuN (neurons) at E11.5 and E12.5 (**Fig. 2D, E**). We counted the number of cycling cells and neurons present, to generate a ratio of neurons to cycling cells in both the control and mutant. We found a significant increase in the ratio of NeuN+/Ki67^+^ cells at E12.5, but not at E11.5 (**Fig. 2D, E**). Altogether, minor spliceosome inhibition reduced VD progenitor numbers and increased neuron production.

### *Rnu11*-null VD resulted in gross dysgenesis of hypothalamic architecture

Given the developmental defects in the *Rnu11*-null VD, we next looked at the gross morphology of the hypothalamus. Towards this end, we collected serial coronal sections of P30 brains to evaluate the tdTomato footprint, followed by Nissl staining to assay the gross histological organization of the hypothalamus. Representative white light and tdTomato images across the hypothalamus revealed disruption in the structure of the tuberal and posterior regions of the hypothalamus in the mutant (**Fig. 3A**). Notably, the third ventricle, which is present and observable in the anterior hypothalamus, does not extend to the tuberal and posterior hypothalamus of mutant mice (**Fig. 3A**). Furthermore, Nissl analysis revealed the loss of discernable nuclei, including the DMH, VMH, and ARC, within the tuberal hypothalamus (**Fig. 3A**). Despite the structural deficits we observed in the mutant hypothalamus by Nissl staining, 3D reconstruction of the hypothalamic tdTomato+ domain showed that the mutant VD manages to produce a hypothalamus-like structure, albeit smaller than the control (**Fig. 3B,B’**).

**Figure 3.**
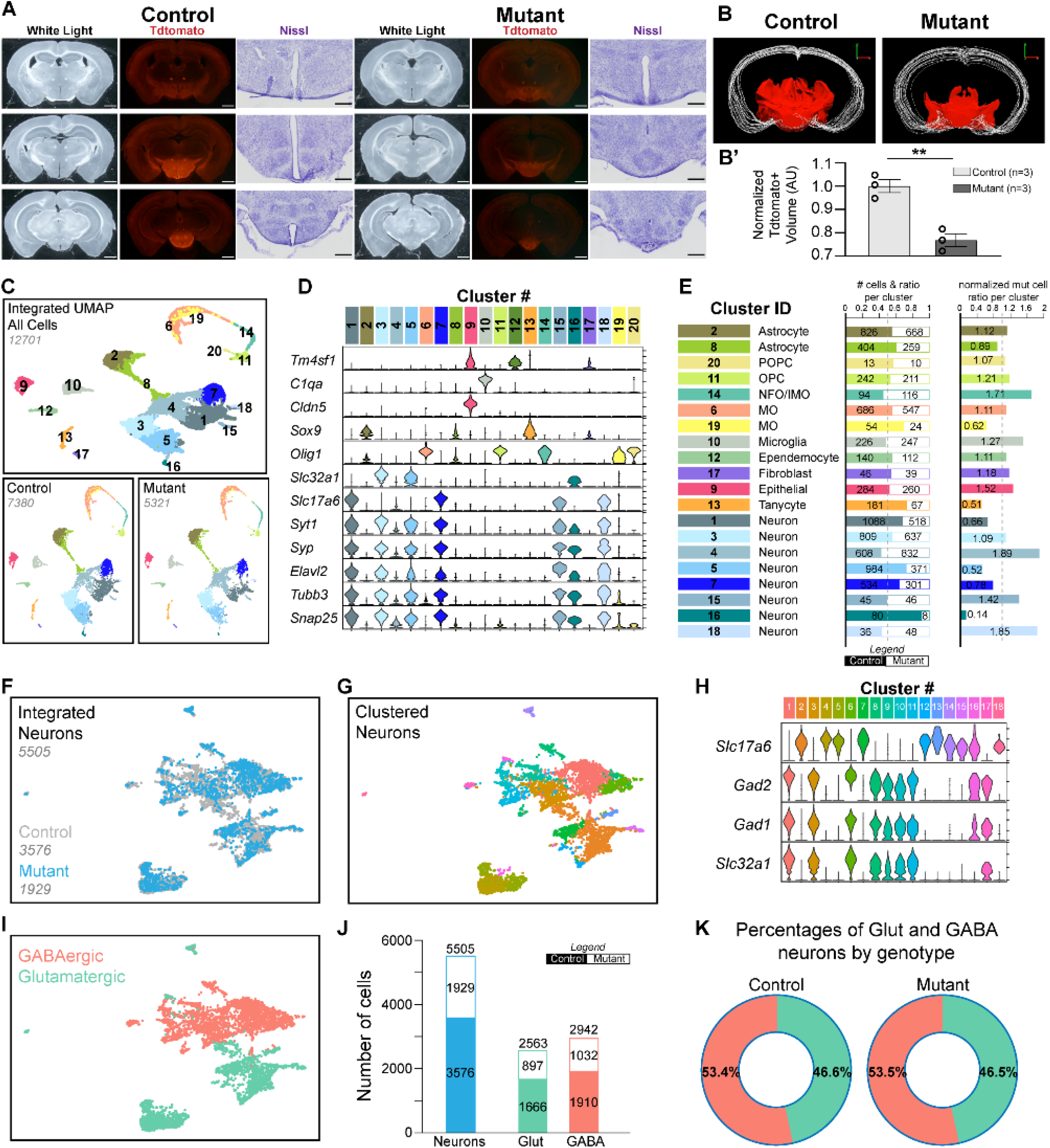
All major cell types present despite the reduced hypothalamic size in *Rnu11* cKO mouse. White light and fluorescent tdTomato-reporter images of control (left) and mutant (right) P30 female brains in the rostral (top) to caudal (bottom) axis. Images of the hypothalamus of the same sections after Nissl staining. 3D reconstruction of *Nkx2*.*1-Cre*+ lineage on juvenile control (left) and mutant (right) brains. (**b**’) Quantification of normalized *Nkx2*.*1-Cre+* lineage hypothalamic to brain volume in control (light grey, n=3) and mutant (dark grey, n=3) mice. (**c**) Top, unsupervised clustering of all, high quality, integrated cells from control and mutant hypothalami represented in a uniform manifold approximation projection (UMAP), colored coded by cluster. Bottom, UMAP of all cells color in control (left) and mutant (right) cells, color coded by cluster. (**d**) Violin plot of the pan-neuronal markers (*Snap25, Tubb3, Elavl2, Syp, Syt1*), neurotransmitter components (*Slc32a1 and Slc17a6*) and non-neuronal discriminatory markers (*Olig1, Sox9, Cldn5, C1qa, Tm4sf1*). (**e**) Cluster identification based on top genes expressed within each cluster. (**e’**) Number and ratio of the contribution of control and mutant per cluster. Dotted grey line at 0.5 represents where the number of control and mutant cells per cluster are equal. (**e’’**) Ratio of mutant cells per cluster to control cells per cluster, normalized to the input number of control and mutant captured cells. Dotted grey line at 1 represents where the proportion of mutant cells in a cluster over the input mutant cells is equivalent to the number of control cells per cluster over the input number of control cells. (**f**) UMAP of integrated neurons colored coded by genotype, control (grey) and mutant (blue). (**g**) UMAP of integrated neurons colored coded by unsupervised clustering analysis. (**h**) Violin plot, in each cluster, GABAergic (*Slc32a1, Gad1, Gad2*) and Glutamatergic (*Slc17a6*) neurotransmitter components/enzymes of the 18 neuronal clusters. (**i**) UMAP of integrated neurons colored GABAergic and Glutamatergic. (**j**) Bar graph of total neurons (blue), and breakdown of GABAergic (salmon) and glutamatergic (teal), with control (filled bar) and mutant (empty bar). (**k**) Donut graph of the percentage of GABAergic and glutamatergic neurons out of all neurons, by genotype. Data (**b’**) represented as mean, with error bars ±SEM. Significance was determined by two-tailed student’s t-tests. **=*P*<0.01 (See also **Supplementary Data Table 9**).

### The mutant hypothalamus contains all key neuronal and non-neuronal cell types

The gross morphological alterations in structure of the mutant hypothalamus led us to investigate the cellular composition of the hypothalamus in mutant mice. To assess the cell type composition, we microdissected the tdTomato+ hypothalamic footprint and performed scRNAseq on control and mutant female juvenile mice (P30; *N*=2, with each *N* containing 3 pooled hypothalami) (**Fig. 3A, Supplementary Data 2A**). After filtering cells with >40% mitochondrial reads or less than 500 unique molecular identifiers (UMIs), we obtained 7380 high-quality control cells and 5321 high-quality mutant cells (**Fig. 3C**). In the control data set, the median number of genes per cell were 2538.5, and median UMIs per cell was 5507 (**Supplementary Data 2B**). In the mutant data set, the median number of genes per cell were 2385, and median UMIs per cell was 4906 (**Supplementary Data 2B**). Comparing the top 10 discriminatory genes in the control and mutant scRNAseq data showed distinct patterns. While there was some overlap (*Avp, Gal, Sst*), the mutant hypothalamus lacked key neuropeptides regulating feeding (*Agrp, Pomc, Npy*) (**Supplementary Data 2C**).

To ensure that the same filtering criteria and gene sets were driving cell population identification and clustering, irrespective of the genotype, we integrated the control and mutant scRNAseq data. Through unsupervised clustering of all the integrated cells, we generated a uniform manifold approximation projection (UMAP) and identified 20 clusters (**Fig. 3C**). The mutant hypothalamus was smaller (**Fig. 3A-C**) and fewer cells were captured (**Fig. 3C**). Despite this, all the major cell clusters identified in the control hypothalamus, were also found in the mutant (**Fig. 3C**). We used the aggregated expression of known pan-neuronal markers (*Snap25, Syp, Tubb3, Elavl2)* (**Fig. 3D**) to identify 7 neuronal clusters (1, 3, 5, 7, 15, 16, and 18) and 13 non-neuronal clusters (2, 4, 6, 8, 9, 10, 11, 12, 13, 14, 17, 19, and 20) (**Fig. 3D**). Based on the top, highly expressed genes as well as previously published hypothalamic scRNAseq data^26,27^, we identified 2 astrocyte populations, 5 populations within the oligodendrocyte lineage, 1 microglia population, as well as single epithelial, tanycyte, ependemocyte, and fibroblast populations (**Fig. 3E**). Broadly, non-neuronal populations were both identified and segregated out by high expression of markers including *Tm4sf1* (Clusters 9, 10, 12, 13), *Cldn5* (Cluster 10), *Sox9* (Clusters 2, 8 13, *C1qa* (Cluster 1), and *Olig1* (Clusters 6, 19, 14, 20, 11) (**Fig. 3D, E**).

To assess changes in neuronal populations that may be reflective of the developmental insult, we performed unsupervised clustering of the neurons, to establish 18 neuronal clusters (**Fig. 3F-H**). Consistent with previous results (**Fig. 3C**), separation of these pooled neurons by genotype, identified 3575 control and 1929 mutant neurons (**Fig. 3F**). Next, we used the expression of *Slc32a1, Gad1*, and *Gad2* to identify GABAergic neurons (Clusters 1, 3, 6, 8, 9, 10, 11, and 17) and *Slc17a6* to identify glutamatergic neurons (Clusters 2, 4, 5, 7, 12, 13, 14, 15, 16, and 18) (**Fig. 3H,I**). When sorted by genotype, we found reductions in both GABAergic and glutamatergic neurons (**Fig. 3J**). However, when we normalize to the number of neurons captured, we found the same percentage of GABAergic and glutamatergic neurons in mutant as the control (**Fig. 3K**).

### scRNAseq revealed stark reductions in ARC neuronal subtypes regulating feeding

The distribution of the broad GABAergic and glutamatergic neurons suggested that the mutant hypothalamus had appropriate ratios of the expected neuronal populations. To further probe this idea, we performed supervised, iterative clustering on glutamatergic neurons, to obtain 15 clusters (**Supplementary Data 3, Fig. 4A**). Like the integrated analyses of all cell clusters, we found that there are cells from both control and mutant present within each of the clusters identified (**Fig. 4B, Supplementary Data 3B**). Identification of glutamatergic subtypes within the data set was through the transcriptional profiles of the neurons (**Fig. 4C**). Given the broad hypothalamic dissection we captured many previously documented glutamatergic clusters. For example, Cluster 7, identified by high expression of *Foxb1*, likely corresponded to the mammillary body of the posterior hypothalamus^26,28^(**Fig. 4C**). Further, Cluster 9, identified by co-expression of *Oxt* and *Avp*, likely corresponded to two closely related populations, oxytocin (OXT) and arginine vasopressin (AVP) neurons, respectively, of the PVH and SON, which may also include AVP neurons of the SCN^26,48^ (**Fig. 4C**). Indeed, when we separately examined this Glutamatergic Cluster 9, we further refined that there are two distinct populations (*Oxt*^+^/*Avp*^-^ and *Avp*^+^/*Oxt*^-^, **Supplementary Data 4A**,**B**). *In situ* for *Oxt* and *Avp* showed intertwined localization of these neurons within the PVN and SON in both the control and mutant, with apparent disorganization and reductions in the ratios of these neurons in the SCN (**Supplementary Data 4C**). Importantly, we identify neuronal populations of the LH and ARC, which were the regions with distinct morphological changes in mutant mice (**Fig. 3A**). For example, Cluster 12 corresponds to ARC *Pomc*^+^ satiety signaling neurons and Cluster 11 corresponds to the LH *Hcrt*^+^ neurons, regulating behaviors including arousal, stress, and reward^16,27^ (**Fig. 4C**). Given their presence in the compromised mutant hypothalamus, we wanted to understand the relative shift in the number of these subtypes. Therefore, we first normalized the number of mutant cells in each cluster over total number of mutant glutamatergic cells, which was normalized by the value of the number of control cells in each cluster over total number of control glutamatergic cells. While most of the glutamatergic neuronal subtypes (10 clusters) were underrepresented in the mutant, some neuronal populations are revealed to be unchanged or overrepresented (Clusters 3, 11) in the mutant (**Fig. 4D, Supplementary Data 3B**). Given that the gross morphology of the mutant hypothalamus is severely compromised, we next wanted to study how the scRNAseq data would manifest in 3D. For this, we performed *in situ* hybridization for *Pomc* (Cluster 12) and *Hcrt* (Cluster 11) on serial sections across the control and mutant hypothalami (**Supplementary Data 5D-G**). We then reconstructed this data in 3D to validate the ratiometric shifts identified by the scRNAseq analysis, as well as to gain the spatial resolution of these neuronal subtypes that was lost in the single cell dissociation (**Fig. 4E, Supplementary Data 5E**). We observed that *Pomc*^+^ neurons are severely reduced in the mutant, validating the scRNAseq data, and that *Pomc*^+^ neurons do not span comparably within the rostral to caudal axis (**Figs. 4E,F; Supplementary Data 5G**). Similarly, we show that Hcrt^+^ neurons are only mildly reduced in raw number in the mutant, validating the identified normalized overrepresentation from the scRNAseq data, and that *Hcrt*^+^ neurons were spatially organized in a comparable pattern to the control (**Supplementary Data 3B, 5D-F**).

**Figure 4.**
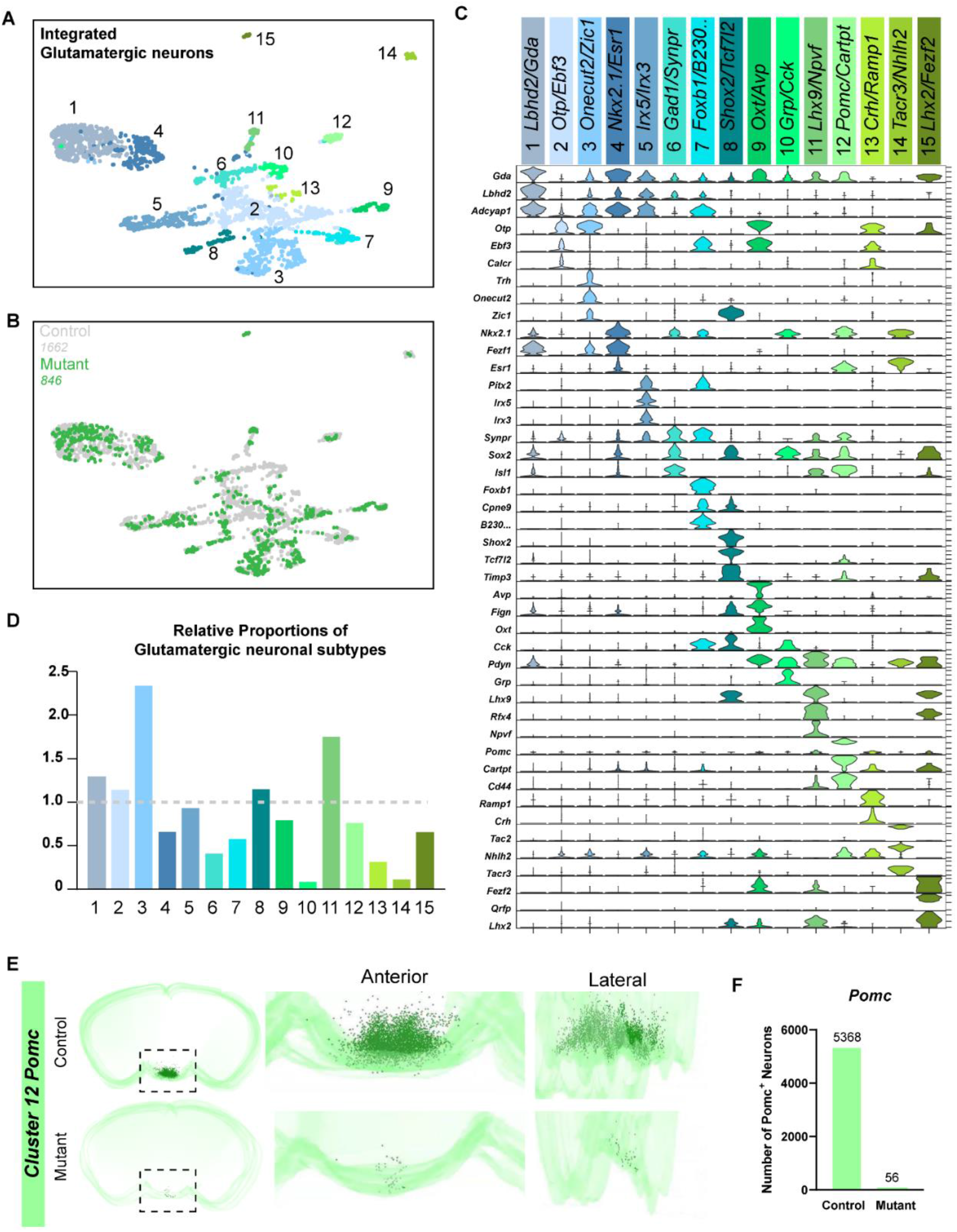
Ratiometric shifts in key glutamatergic neuronal populations integral to feeding, satiety and arousal. (**a**) Supervised clustering of integrated glutamatergic neurons from control and mutant hypothalami represented in a UMAP, colored coded by cluster. (**b**) Bottom, UMAP of integrated glutamatergic neurons, color coded by genotype. Control=grey; Mutant=green. (**c**) Violin plots of the top 3 discriminatory genes per integrated glutamatergic cluster. (**d**) Ratio of the number of mutant glutamatergic cells per cluster to control glutamatergic cells per cluster, normalized to the total number of respective control and mutant glutamatergic neurons captured. Dotted grey line at 1 represents where the proportion of mutant cells in a cluster over the input mutant cells is equivalent to the number of control cells per cluster over the input number of control cells. (**e**) 3D rendering of in situ hybridization results for *Pomc* in P30 juvenile control and mutant. Zoomed inset of 3D reconstruction from anterior and lateral views. *Pomc*^+^ neurons = green dots. (**f**) Quantification of *Pomc* neurons in control and mutant P30 hypothalami (n=1, serial sections across full hypothalamus).

Iterative, supervised clustering of the GABAergic neurons resulted in 14 GABAergic and 1 histaminergic cluster (**Supplementary Data 3A)**. The emergence of this histaminergic cluster from the GABAergic neurons was driven by high expression of *Hdc*, encoding histidine decarboxylase the enzyme necessary for production of histamine^49^ (**Supplementary Data 3A**). This cluster of 16 total cells, also co-expressed previously reported key markers including *Itm2a, Maob, Slc18a2*, and *Prph*^26,28,31^ (**Supplementary Data 3A**,**D**). Of the 16 identified histaminergic neurons, only 2 were from the mutant. Serial section *in situ* hybridization analysis for *Hdc* showed reduced numbers, yet comparable spatial distribution in the mutant hypothalamus (**Supplementary Data 3A**,**D, 5A-C**).

Of 14 GABAergic clusters (**Supplementary Data 3, Fig. 5**), 13 consisted of both control and mutant neurons, with the exception of Cluster 11 (*Sox14*/*Unc13c*), which lacked mutant neurons (**Fig. 5A, B**). Consistent with previous analyses^26–28^, we identified many known GABAergic subpopulations across the hypothalamus (**Fig. 5C**). For example, Cluster 6 (*Crabp1/Tbx3*), corresponds to a reported neuronal population of the ARC^26^. Moreover, Cluster 10 (*Ptk2b/Pthlh*), likely represents a GABAergic population of the LH^27,28^. To determine the ratiometric distribution of the identified GABAergic neuronal subtypes, we then applied the aforementioned normalization. The majority (11 clusters) were underrepresented in the mutant compared to the control, while some were unchanged (Clusters 4, 10, 12) and overrepresented (Clusters 1, 5, 13) (**Fig. 5D**). Of note, Cluster 9 (*Agrp+/Npy+*) was drastically underrepresented in the mutant relative to the control (**Fig. 5D**). Given that *Npy*+ neurons act antagonistically to *Pomc*^+^ neurons, which were also reduced (**Fig. 4E,G**), we used *in situ* hybridization for *Npy* to determine the spatial resolution. Indeed, we observed severe reduction in *Npy*^+^ neurons in the mutant hypothalamus, with clear staining for *Npy*^+^ interneurons within the cortex of the same section, showing that this is hypothalamus-specific (**Fig. 5E, E’**). Given the stark reduction in both GABAergic and glutamatergic neuronal populations, especially those within the ARC regulating feeding and satiety, we hypothesized that these mice would have impaired energy balance, including feeding, postnatally.

**Figure 5.**
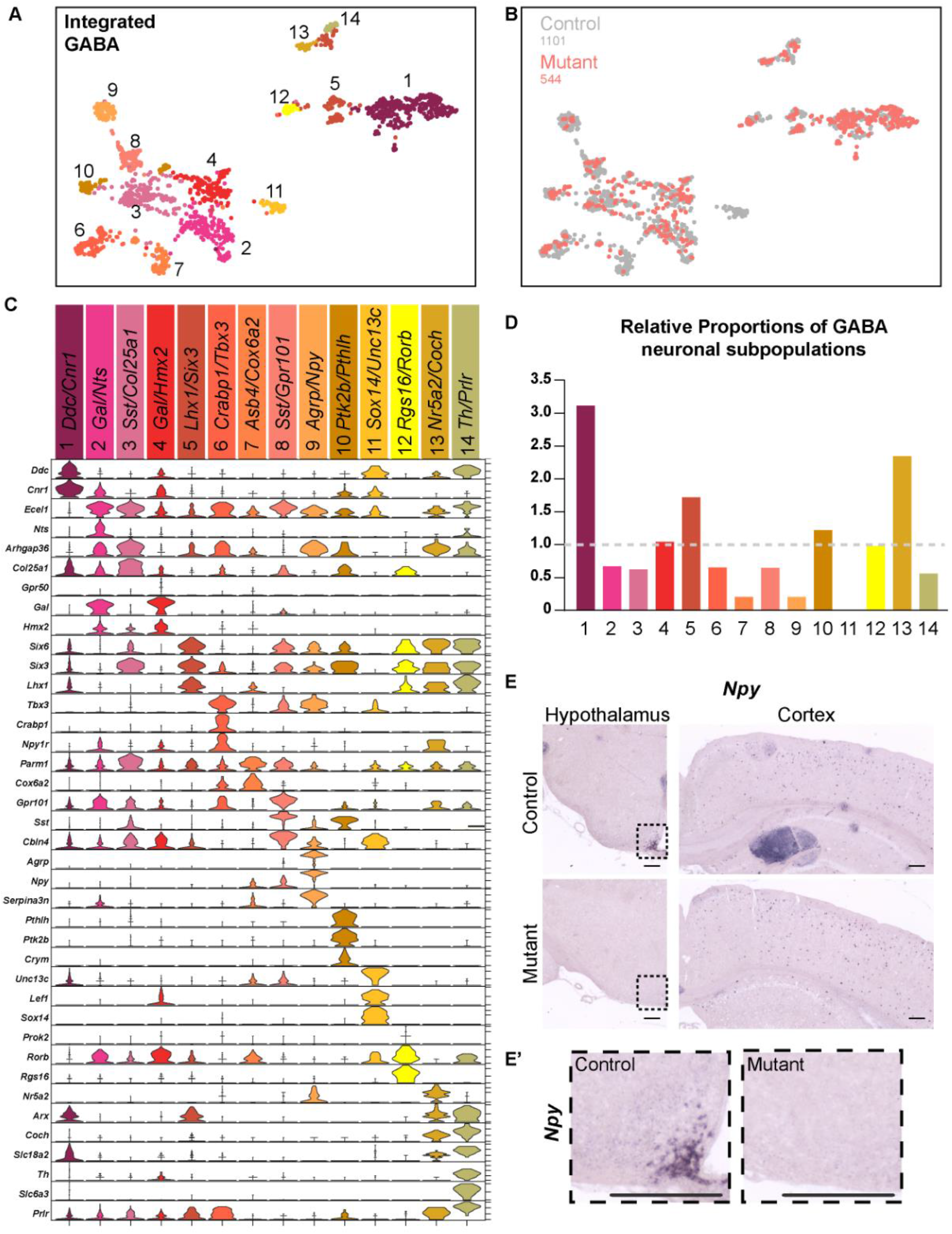
Broad reductions in the proportions of GABAergic neuronal populations in the *Rnu11*-null hypothalamus. (**a**) Supervised clustering of integrated GABAergic neurons from control and mutant hypothalami represented in a UMAP, colored coded by cluster. (**b**) Bottom, UMAP of integrated GABAergic neurons, color coded by genotype. Control=grey; Mutant=salmon. (**c**) Violin plots of the top 3 discriminatory genes per integrated GABAergic cluster. (**d**) Ratio of the number of mutant GABAergic cells per cluster to control GABAergic cells per cluster, normalized to the total number of respective control and mutant GABAergic neurons captured. Dotted grey line at ratio of 1. (**e**) *In situ* hybridization for *Npy* in the arcuate nucleus of the hypothalamus (left) and the cortex (right), in control (top) and mutant (bottom) P30 juvenile brains. (**e’**) Magnified image of the arcuate showing *Npy* signal in the control(left) and mutant (right). Scale bar 200µm.

### Ablation of *Rnu11* in the developing VD results in failure to thrive, and subsequent obesity

To determine in if ablation of *Rnu11* in the developing VD impaired energy balance, we first monitored postnatal development by measuring body weight of control and mutant mice. At birth, mutant mice were indistinguishable from controls, but showed delayed growth by P7 (**Fig. 6A-B, D,E**). At weaning (P21), mutants were smaller than controls (**Supplementary Data 6A,B**), and the shift to *ad libitum* food, was followed by rapid weight gain and obesity (**Fig. 6C,F**). Given that postnatal weight regulation is highly sex-dependent, we analyzed body weight as a function of sex and age. We found that, compared to controls, female mutants were initially underweight through 4 weeks of age, at an equal weight at 5 weeks of age, and were significantly overweight at 7 weeks of age (**Fig. 6G**). In contrast, male mutant mice showed a delayed crossover in weight (at 7-8 weeks) and significantly outweighed controls by 13 weeks of age (**Fig. 6H**). Comparing slopes of weight kinetics between the control males and females revealed that males exhibit a significantly higher rate of weight gain than females (**Supplementary Data 6C**), which was not observed in the mutant animals (**Supplementary Data 6D**). Since female mutants exhibited early and robust susceptibility to weight gain, subsequent metabolic and feeding analysis was focused on characterizing this female phenotype.

**Figure 6.**
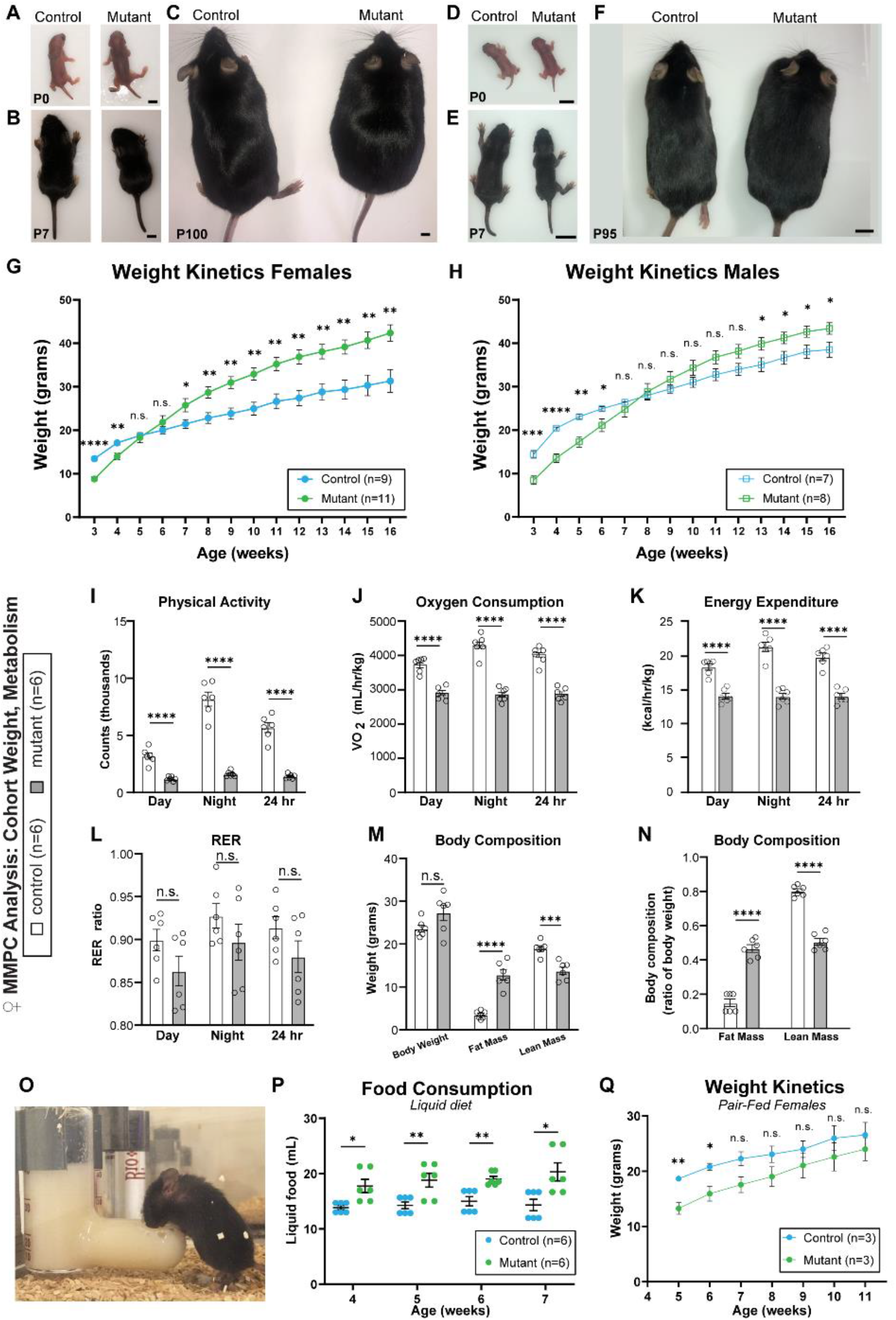
Early failure to thrive is followed by obesity, due to hyperphagia in the *Nkx2*.*1-Cr*e *Rnu11* cKO mouse. (**a**) Comparable body size in control (left) and mutant (right) P0 female mice. (**b**) Reduced size of mutant female mouse (right) at P7, relative to control (left). (**c**) Visible obesity in the mutant female at P95. (**d-f**) Postnatal progression of male mice at P0 (**d**), P7(**e**), and P100 (**f**). (**g-h**) Weight kinetics of female (**g**) and male (**h**) control (blue) and mutant (green) mice. (**i**) Physical activity as measured by counts of beam break across day, night and over 24-hour period in control (white) and mutant (grey) mice. (**j**) Oxygen consumption of control (white) and mutant (grey) mice during day, night, and across total 24-hour period represented as the volume of oxygen consumed in mL/hr/kg. (**k**) Calculated energy expenditure rate (kcal/hr/kg) in control (white) and mutant (grey).(**l**) Respiratory exchange ratio of control (white) and mutant (grey) mice during day, night, and across total 24-hour period. (**m)** Body weight, fat mass and lean mass measurements in grams of control (n=6, white) and mutant (n=6, grey) female mice. (**n**) Body composition normalized to percent of body weight of fat mass and lean mass in control (white) and mutant (grey) mice. (**o**) Image of mutant female mouse accessing liquid food diet in L-shaped food dispenser (Bioserv. #9019, F1268SP). (**p**) Bar graph displaying the average daily liquid food consumption averaged weekly by female control (blue) and mutant (green) mice through 7 weeks of age. (**q**) Weight kinetics of pair-fed control (blue, n=3) and mutant (green, n=3) female mice. Data (**g-n, p, q**) represented as mean, with error bars ±SEM. Significance was determined by two-tailed student’s t-tests. n.s.= not significant, *=*P*<0.05, **=*P*<0.01, ***=*P*<0.001, ****=*P*<0.0001 (See also **Supplementary Data Table 9**).

### Metabolic defects exacerbate the obesity observed in *Rnu11*-null mice

To determine how alterations in energy balance contribute to obesity following ablation of *Rnu11* in the developing VD, control and mutant female mice were subjected to metabolic and body mass analyses. We observed a suppression of physical activity (**Fig. 6I**), whole-body oxygen consumption (**Fig. 6J**), and energy expenditure (**Fig. 6K**) in mutant females across the circadian light cycle. A similar respiratory exchange (approaching 1) was observed between control and mutant mice, suggesting both groups used carbohydrates as the primary energy substrate for oxidative metabolism (**Fig. 6L**). Importantly, prior to the metabolic analysis, using *in vivo* whole-body proton magnetic resonance imaging, we confirmed that mutant females were weight matched in body weight to controls, ruling out the possibility that weight changes accounted for differences in energy balance of mutant mice (**Fig. 6M**). Nonetheless, mutant females had increased proportion of fat to lean body mass (50:50) compared to control mice (20:80) (**Fig. 6N**), confirming that ablation of *Rnu11* in the developing VD promotes obesity by increasing body fat in adulthood. Altogether this analysis revealed that obesity in adult mutant mice is in part due to disruptions in energy expenditure.

Given that *Nkx2*.*1-Cre* has been shown to drive recombinase expression in cells of the VD, as wells as the pituitary and thyroid^8,9^, we assayed hormones of the hypothalamus-pituitary-thyroid-axis in the same cohort of weight-matched mice. Serum levels of adrenocorticotropic hormone (ACTH) and prolactin (PRL) were comparable between controls and mutants (**Supplementary Data 7A**). However, consistent with the short stature observed in mutant mice (**Supplementary Data 6A**,**B**), growth hormone (GH) was significantly reduced in the mutants (**Supplementary Data 7A**). This GH-specific phenotype was observed despite the apparent global reduction in total pituitary size in the mutant compared to the control (**Supplementary Data 7B**). Furthermore, analysis of hormones within the HPT-axis revealed a significant increase in thyroid stimulating hormone (TSH) concurrent with a significant decrease in thyroxine (T4), with comparable levels of thyrotropin releasing hormone (TRH) (**Supplementary Data 7A**). This suggests a lack of the negative feedback loop from the pituitary in response to sufficient levels of thyroid hormone and is reflective of the decreased thyroid size (**Supplementary Data 7C**). Taken together, the mutant mice exhibit both metabolic dysfunction and decreased energy expenditure.

### Hyperphagia drives obesity transition in *Rnu11* cKO *Nkx2*.*1-Cre*+ mice

Weight gain is ultimately the result of interactions between caloric intake, energy expenditure, and metabolism. To begin disentangling these variables, we next explored energy consumption by employing a liquid food consumption assay in the home cages of control and mutant female mice (**Fig. 6O**). Starting at 4 weeks of age, mutant mice consumed significantly more food compared to controls, which continued throughout the experiment (**Fig. 6P,G**). Thus, the mutant mice had elevated energy consumption, combined with reduced energy expenditure (**Fig. 6I-N**). Therefore, we sought to decouple the primary initiator of the obesity phenotype by pair-feeding assay in which mutant mice were fed the amount of food consumed by age-matched controls. Across the 11 weeks of experimentation, mutant pair-fed mice remained significantly underweight through 6 weeks and did not surpass controls (**Fig. 6Q**).

To further understand the specific pattern of hyperphagia in these mice we implemented the Feeding Experimentation Device 3 (FED3) to reliably monitor free feeding in these mice across the light:dark cycles^50^ (**Fig. 7A**). Consistent with liquid food diet analysis, we found that mutant mice retrieved significantly more pellets than controls (**Fig. 7B**). Unlike the controls, which retrieve significantly more pellets in the dark cycle, the mutant mice, retrieved comparable pellets across both the dark and light cycles (**Fig. 7B**). Specifically, the total pellets retrieved by the mutant in the light cycle is significantly more that control in the light cycle, which is not the case for the number of pellets retrieved in the dark cycle (**Fig. 7B**). Given that mutant mice consumed more food, we next analyzed the median meal size of controls and mutants, which revealed that control mice tended to have more frequent 1 pellet meals, whereas mutants prefer 2-3 pellets per meal (**Fig. 7C**). On average, however, mutants had a significantly higher count of 3 pellet meals, while all other meal sizes were comparable between the controls and mutants (**Fig. 7C**).

**Figure 7.**
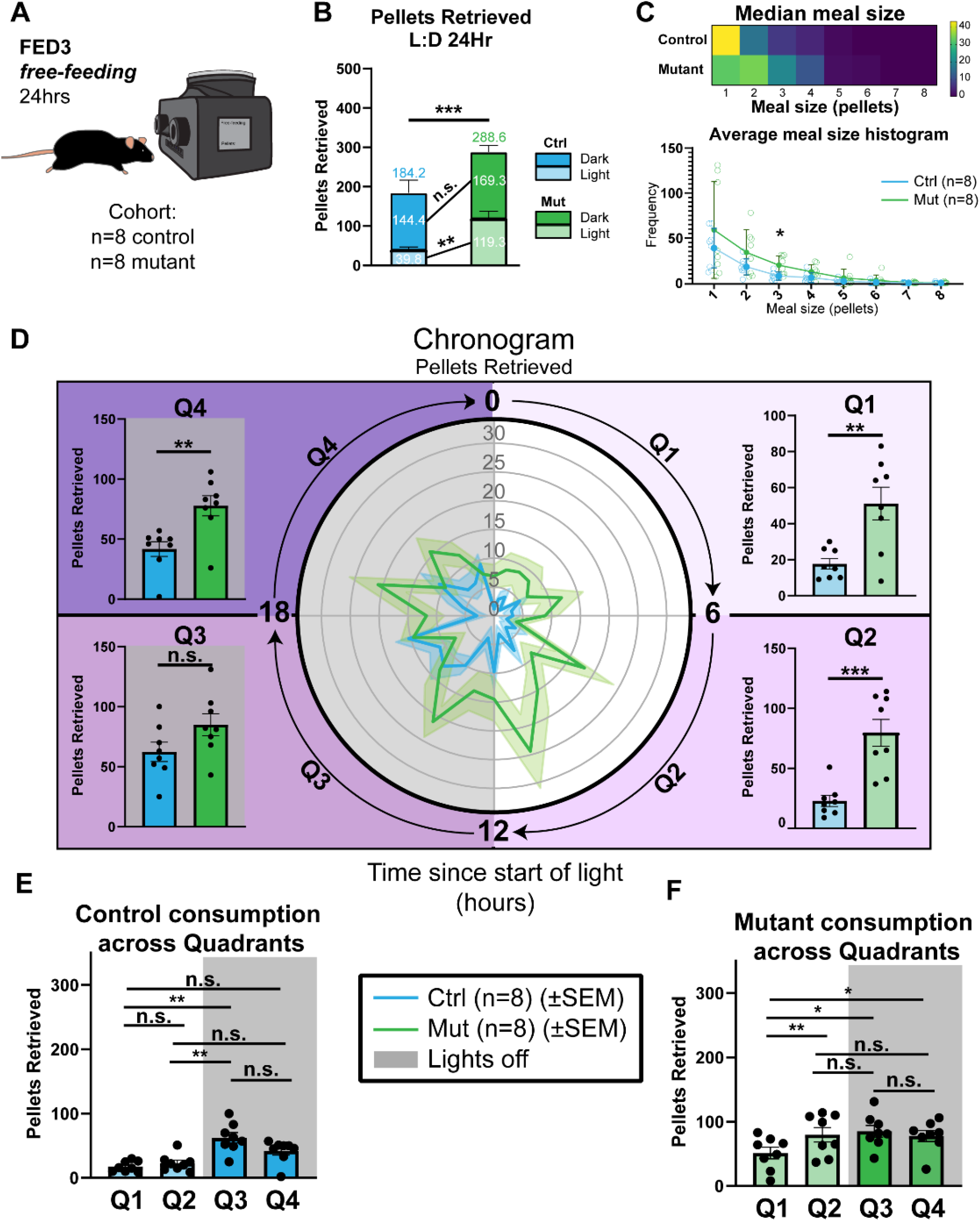
Satiety and circadian feeding patterns altered in *Rnu11*-null mutant mice. Schematic representation of Feeding Experimentation Device version 3 (FED3) in free-feeding mode and cohort of females. (**b**) Stacked bar chart of pellets retrieved in a 24hr period by control (blue) and mutant (green) mice across the light (opaque) and dark (full color) cycles. (**c**) Meal analysis of control and mutant mice. Top, heatmap of median meal sizes of control and mutant mice during the 24hr period. Bottom, average meal size histogram of control (left, blue) and mutant (right, green) across the recording period. (d) Chronogram of pellets retrieved by control (blue) and mutant (green) mice. Radial circles from center quantify average number of pellets consumed and clockwise rotation around the circle denotes time in hours from start of the light phase (0) to start of the dark phase (12) back to start of the light phase (0). Bar graphs of food consumption broken down by quadrant (Q) between control and mutants. Q1 encompassing hours 0-5 of light cycle (top right, light lavender), Q2 encompassing hours 6-11 of light cycle (bottom right, lavender), Q3 encompassing hours 12-17 of light cycle (bottom left, purple), and Q4 (hours 18-23 of light cycle (top left, dark purple). (**e**) Pellets retrieved by the control across Q1-Q4. (**f**) Pellets retrieved by the mutant across Q1-Q4. Data (**b, c**(bottom), **d**) represented as mean, with error bars ±SEM. Significance was determined by two-tailed student’s t-tests. n.s.= not significant, *=*P*<0.05, **=*P*<0.01, ***=*P*<0.001. Data (**e, f**) represented as mean, with error bars ±SEM. Significance was determined by ANOVA, followed by post-hoc Tukey’s test. n.s.= not significant, *=*P*<0.05, **=*P*<0.01, ***=*P*<0.001 (See also **Supplementary Data Table 9**).

A chronogram of pellet retrieval across a 24hr period revealed that mutant mice consistently consumed food across both the dark and light cycles (**Fig. 7D**). Analysis of pellet retrieval in bins of 6-hour increments, showed that the mutant mice consume significantly more food in 3 of the 4 quadrants (Qs), with the exception of Q3, which marks the beginning of the dark cycle when control mice retrieve the largest number of pellets (**Fig. 7D,E**). When we compare pellet retrieval by the mutant across the 4 quadrants, we found that mutant mice eat significantly less food in Q1 (**Fig. 7F**), but mutants still eat more in Q1 than controls (**Fig. 7D**). Ultimately, we showed that ablation of *Rnu11* in the *Nkx2*.*1*+ domain promoted hyperphagia across the circadian day, driving obesity.

## Discussion

Here, we show that integrity of the minor spliceosome is essential for hypothalamic development and function. Specifically, we show that ablation of *Rnu11* inhibits the minor spliceosome, which resulted in elevated minor intron retention and aberrant AS of MIGs involved in cell cycle regulation, consistent with previous reports^40,51,44,43^. As expected, we found progenitor cell cycle defect in the VD (**Fig. 1G,H**), that ultimately resulted in cell death (**Fig. 1I,J**). The developing VD responds to the progenitor cell loss by proportionally reducing the various progenitor subdomains (**Fig. 2A**). Although *Shh* is not a marker of VD progenitors, it marks the boundary of the VD and the shift in its expression profile in the mutant further illustrates the response of the developing hypothalamus to progenitor cell loss (**Fig. 2A**). This plasticity of the developing VD is further illustrated by the unexpected enrichment of the neuronal GOTerms (e.g. “synapse”) (**Fig. 2C**) by the upregulated genes in the mutant VD. This molecular finding was confirmed by the increased ratio of neurons/progenitor cells in the mutant the developing pre-hypothalamus (**Fig. 2C-E**). This finding suggests that upon loss of progenitor cells, a developing system reconfigures its developmental trajectory to approximate the final structure it was originally programmed to generate.

These adaptive changes in the developing hypothalamus were reflected in the P30 mutant hypothalamus, which was smaller than the control (**Fig. 3B**). Despite the reduced size of the mutant hypothalamus, it appeared to produce a structure resembling the control hypothalamus, albeit exhibiting structural abnormalities including the loss of the third ventricle and collapsed tuberal nuclei (**Fig. 3A,B**). Moreover, scRNAseq revealed that the mutant hypothalamus consisted of all the neuronal and non-neuronal cell types observed in the control hypothalamus (**Fig. 3C**). This finding bolsters the idea that the reconfiguration of progenitor subtypes in the developing mutant VD (**Fig. 2A**) allowed the production of a hypothalamus with all its major cell populations (**Fig. 3C**). The varying birth order kinetics of the different neuronal subtypes, combined with progenitor cell loss in the mutant VD, would invariably alter the relative neuronal subtype composition in the adult hypothalamus. Surprisingly, the percentages of GABAergic and glutamatergic neurons were identical between the control and mutant hypothalamus (**Fig. 3K**). This finding suggests that there exists a central organizational principle which ensures balanced production of GABAergic and Glutamatergic neurons. This balance in excitatory to inhibitory ratio could be achieved either during embryonic development or through terminal cell fate shifts. However, amongst the 14 glutamatergic, 15 GABAergic and 1 histaminergic neuronal subtypes identified, we observed clusters that were overrepresented, underrepresented and unchanged in the mutant relative to the control (**Fig. 4, 5 Supplementary Data 3**). Specifically, *Pomc+* (satiety) and *Npy+* (hunger) neurons of the ARC are reduced in the mutant hypothalamus (**Fig. 4, 5**), while *Hcrt+* neurons of the LH were relatively unaltered (**Fig. 4, Supplementary Data 5D-F**).

We show that developmental defects caused by disruption in minor intron splicing compromised the structure and neuronal subtype composition of the hypothalamus, but we also considered the other tissues targeted by *Nkx2*.*1-Cre*, namely the thyroid and pituitary. Both structures were compromised and resulted in metabolic dysfunction, including low GH and T4, which most likely contributed to the pre-weaning failure to thrive, and the weight gain post-weaning (**Supplementary Data 7**). However, pair-feeding experiments allowed us to determine that hyperphagia drove the initial weight gain and progression to obesity, while reduced energy expenditure and physical activity exacerbated the phenotype (**Fig. 6I, K, Q**). Interestingly, analysis of food consumption across 24 hrs revealed that the mutant mice did not eat more at the start of the dark cycle (Q3), when the control eat most of their food (**Fig. 7D,E**). Instead, the mutant mice continue to eat a sustained amount of food across the dark and light cycle (**Fig. 7D,F**).. There are multiple possibilities to explain this hyperphagia. First, the mutant mice are insatiable, which would agree with the fact that they increase their meal size (**Fig. 7C**) and exhibit a reduction in the number of satiety related *Pomc*+ neurons (**Fig. 4**). Second, these mice appear to exhibit disrupted circadian rhythm, and continue to eat across the light cycle, perhaps reflective of the overrepresentation of wake-promoting *Hcrt*+ neurons^16^ (**Fig. 4, Supplementary Data 5D-F**), or disruption of *Nkx2*.*1*-derived circadian, pacemaker SCN neurons which signal to the PVN to inhibit food consumption in response to light^17,48,52^.

Overall, we present that two independent developmental defects (*Rnu11*^FLX/FLX^;*Nkx2*.*1-Cre+* mouse model and PWS), converge in their postnatal phenotypic manifestations (**Fig. 8A**). The combined phenotypes observed in this mouse (pre-weaning failure to thrive, post-weaning hyperphagia, rapid weight gain, obesity, and metabolic dysfunction) overlaps with the symptoms observed in PWS patients^41^(**Fig. 8B**). Moreover, the energy imbalances observed in mutant mice and PWS patients may be driven by similar shifts in neuronal subtype composition as revealed by our scRNAseq and analysis of post-mortem brains of PWS patients^53^ (**Fig. 8C**). For example, we observed reductions in ARC subtypes (*Npy, Pomc*) and unchanged numbers of *Hcrt* neurons, which was also reported in the hypothalami of PWS patients (**Fig. 8C**). This suggests that perturbations in hypothalamic development might play a larger role than appreciated in the mechanism of disease pathogenesis of PWS.

**Figure 8.**
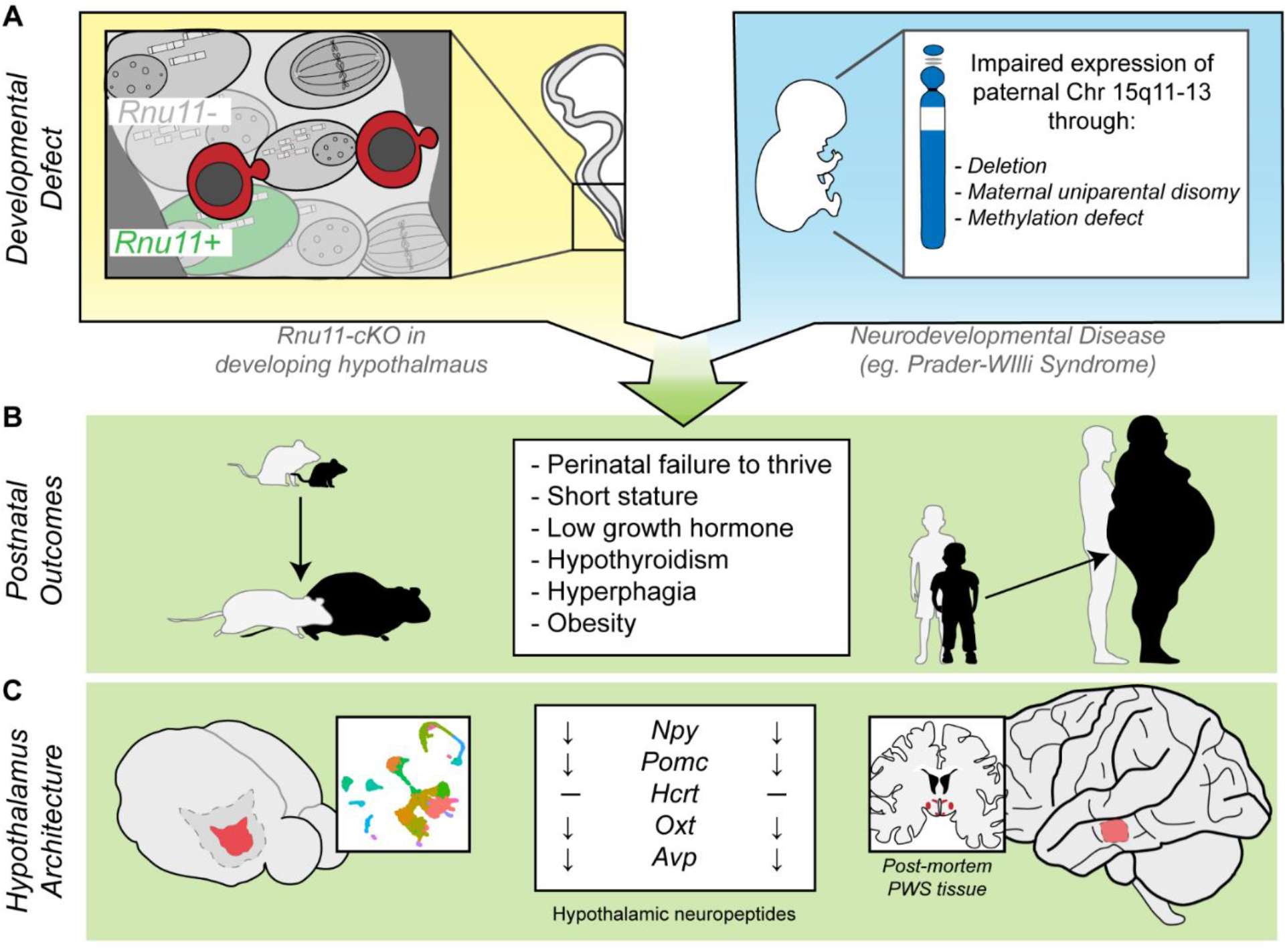
Gross phenotypic and hypothalamic architectural parallels between *Rnu11 Nkx2*.*1-Cre*+ cKO mouse and Prader-Willi Syndrome. (**a-c**) Schematic representations of different developmental defects, leading to shared postnatal outcomes and hypothalamic architecture. (**a**) Left, hypothalamic inhibition of minor intron splicing in mouse, and right, impaired expression of paternal chromosome 15q11-13 in humans as observed in the neurodevelopmental disorder, Prader-Willi Syndrome (PWS). (**b**) Schematic representation of shared postnatal outcomes observed in *Rnu11 Nkx2*.*1-Cre*+ cKO mouse and PWS patients of perinatal failure to thrive followed by hyperphagia driven obesity progression. Center box, list of key phenotypes shared between the mouse model presented and PWS patients.(**c**) Schematic representation of analysis of the hypothalamic architecture in *Rnu11*-null mice and PWS patients. Center box, key hypothalamic neuropeptides underrepresented or unchanged in *Rnu11 Nkx2*.*1-Cre*+ cKO mice (via scRNAseq) and in PWS patients (via post-mortem brain tissue analysis).

## Table of resources/reagents

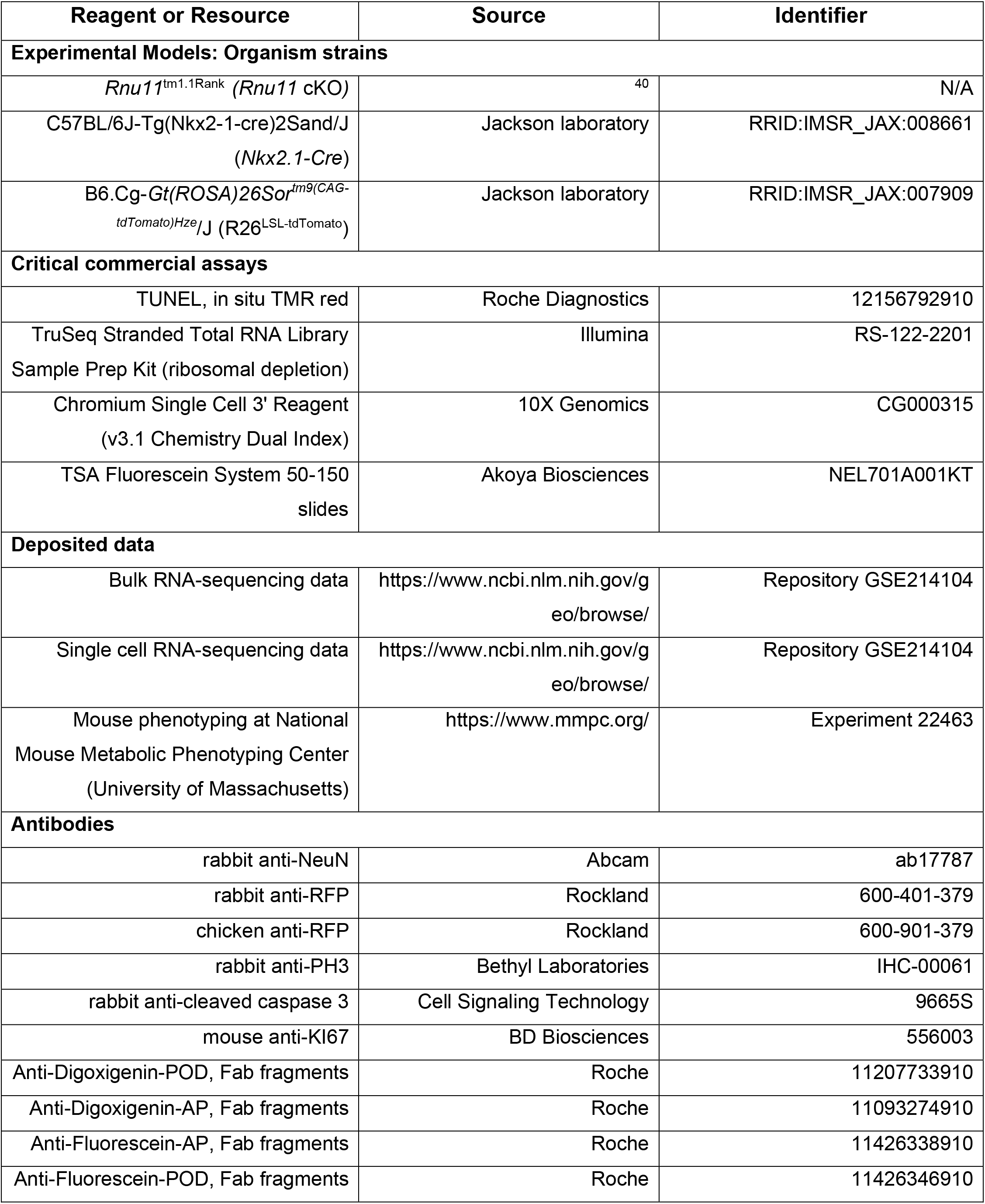
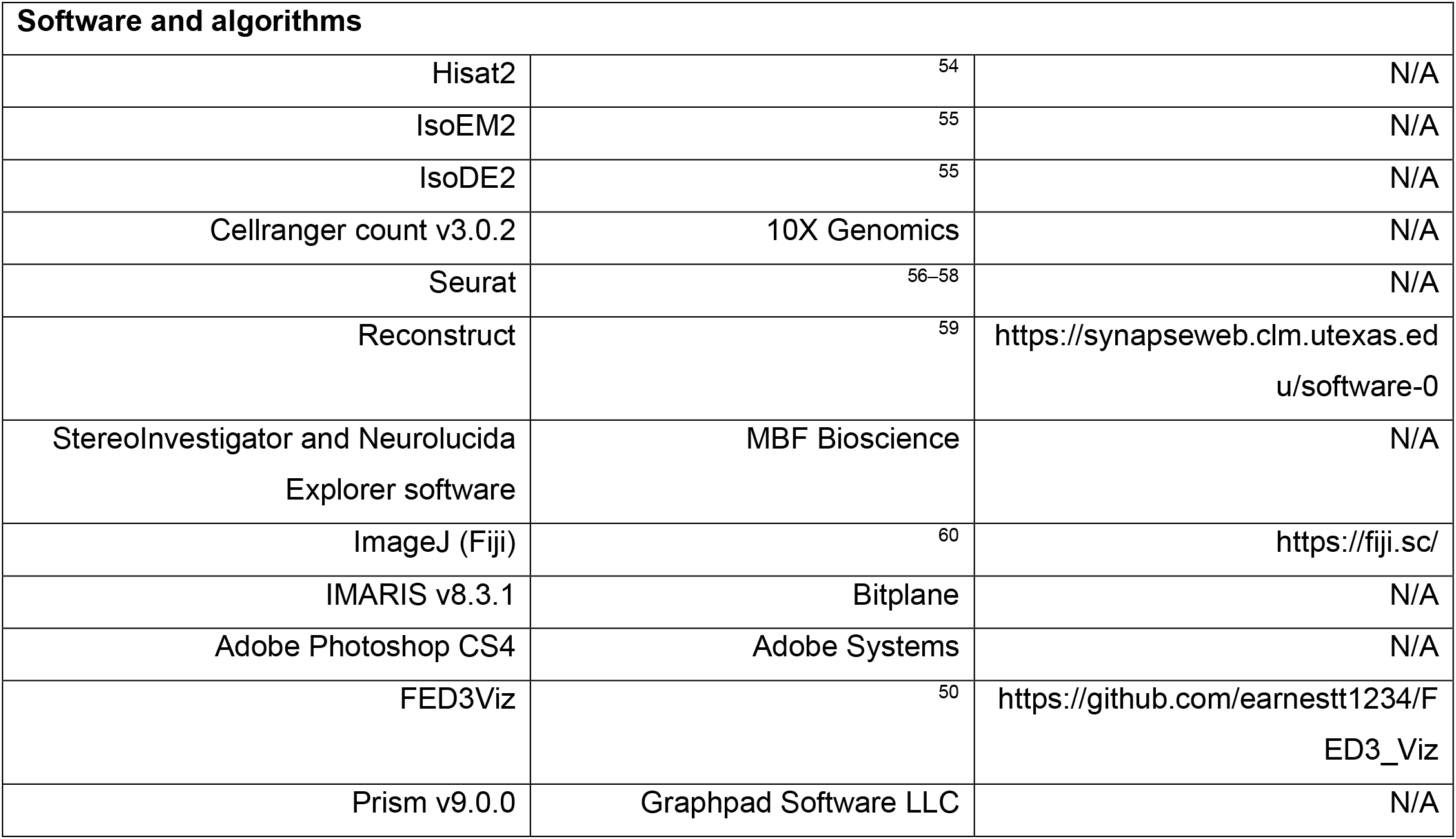

**Table of resources/reagents.**
| Reagent or Resource | Source | Identifier |
| --- | --- | --- |
| **Experimental Models: Organism strains** |  |  |
| *Rnu11*<sup>tm1.1Rank</sup> (*Rnu11* cKO) | 40 | N/A |
| C57BL/6J-Tg(Nkx2-1-cre)2Sand/J<br>(*Nkx2.1-Cre*) | Jackson laboratory | RRID:IMSR_JAX:008661 |
| B6.Cg-Gt(*ROSA*)26Sor<sup>tm9(CAG-tdTomato)Hze</sup>/J (R26<sup>LSL-tdTomato</sup>) | Jackson laboratory | RRID:IMSR_JAX:007909 |
| **Critical commercial assays** |  |  |
| TUNEL, in situ TMR red | Roche Diagnostics | 12156792910 |
| TruSeq Stranded Total RNA Library<br>Sample Prep Kit (ribosomal depletion) | Illumina | RS-122-2201 |
| Chromium Single Cell 3' Reagent<br>(v3.1 Chemistry Dual Index) | 10X Genomics | CG000315 |
| TSA Fluorescein System 50-150<br>slides | Akoya Biosciences | NEL701A001KT |
| **Deposited data** |  |  |
| Bulk RNA-sequencing data | https://www.ncbi.nlm.nih.gov/g<br>eo/browse/ | Repository GSE214104 |
| Single cell RNA-sequencing data | https://www.ncbi.nlm.nih.gov/g<br>eo/browse/ | Repository GSE214104 |
| Mouse phenotyping at National<br>Mouse Metabolic Phenotyping Center<br>(University of Massachusetts) | https://www.mmpc.org/ | Experiment 22463 |
| **Antibodies** |  |  |
| rabbit anti-NeuN | Abcam | ab17787 |
| rabbit anti-RFP | Rockland | 600-401-379 |
| chicken anti-RFP | Rockland | 600-901-379 |
| rabbit anti-PH3 | Bethyl Laboratories | IHC-00061 |
| rabbit anti-cleaved caspase 3 | Cell Signaling Technology | 9665S |
| mouse anti-Ki67 | BD Biosciences | 556003 |
| Anti-Digoxigenin-POD, Fab fragments | Roche | 11207733910 |
| Anti-Digoxigenin-AP, Fab fragments | Roche | 11093274910 |
| Anti-Fluorescein-AP, Fab fragments | Roche | 11426338910 |
| Anti-Fluorescein-POD, Fab fragments | Roche | 11426346910 |

| Software and algorithms |  |  |
| --- | --- | --- |
| Hisat2 | <sup>54</sup> | N/A |
| IsoEM2 | <sup>55</sup> | N/A |
| IsoDE2 | <sup>55</sup> | N/A |
| Cellranger count v3.0.2 | 10X Genomics | N/A |
| Seurat | <sup>56–58</sup> | N/A |
| Reconstruct | <sup>59</sup> | https://synapseweb.clm.utexas.edu/software-0 |
| Stereoinvestigator and Neurolucida Explorer software | MBF Bioscience | N/A |
| ImageJ (Fiji) | <sup>60</sup> | https://fiji.sc/ |
| IMARIS v8.3.1 | Bitplane | N/A |
| Adobe Photoshop CS4 | Adobe Systems | N/A |
| FED3Viz | <sup>50</sup> | https://github.com/earnestt1234/FED3_Viz |
| Prism v9.0.0 | Graphpad Software LLC | N/A |

### Resource availability

#### Lead contact

Further information and requests for resources and reagents should be directed to and will be fulfilled by the lead contact, Rahul Kanadia.

#### Materials availability

This study did not generate new unique reagents.

### Experimental model and subject details

Mouse (*Mus musculus*) husbandry and procedures were carried out in accordance with protocols approved by the University of Connecticut Institutional Animal Care and Use Committee, which operates under the guidelines of the US Public Health Service Policy for laboratory animal care. The *Rnu11* cKO mouse (*Rnu11*^tm1.1Rank^) used in this study was generated and described previously^40^. *Nkx2*.*1*-Cre mouse (Tg(Nkx2-1-cre)2Sand) was bred into the *Rnu11* cKO line to target *Rnu11* for removal in the developing ventral diencephalon^9^. The fluorescent tdTomato reporter mouse (R26^LSL-tdTomato^) was bred into the line to mark both VD progenitors and hypothalamic lineage in the adult^42^. Genotyping for alleles was performed using primers in **Supplementary Data Table 8**. For matings intended for embryonic harvests, E0.5 was considered noon the morning a vaginal plug was observed. The experiments described *Rnu11*^WT/Flx^:: R26^LSL-tdTomato^; *Nkx2*.*1*-Cre^+^ (control), *Rnu11*^Flx/Flx^:: R26^LSL-tdTomato^; *Nkx2*.*1*-Cre^+^ (mutant) embryos and mice, sex denoted per experiment.

## Method Details

### *In situ* Hybridization (ISH)

Cryosections (16 µm) of sagittal embryos (E10.5-E12.5) and coronal whole brains (for P30) (*n*=3 for each timepoint and genotype) were used for section ISH. Section ISH was performed using an antisense, digoxigenin-labeled U11 RNA probe, which was detected using either alkaline phosphatase or fluorescent labeling (FISH), as described previously^40,61^. Primers sequences used for both embryonic and adult *in situs* listed in **Supplementary Data Table 8**.

### TUNEL Cell death assay

Terminal deoxynucleotidyl dUTP transferase nick end labeling (TUNEL) was performed on 16 µm coronal cryosections across the ventral diencephalon (E10.5-E12.5) using the *in situ* cell death detection kit, TMR Red (Roche Diagnostics, 12156792910), in accordance with the manufacturer’s instructions.

### Bulk RNAseq of embryonic diencephalon

TdTomato+ ventral diencephalon was micro dissected under UV scope from control (n=3) and mutant (n=3) E12.5 embryos. Tissue was suspended in 200uL of Trizol and total RNA was extracted using phenol-chloroform extraction, including DNAase treatment. Library sample preparation and sequencing were executed by the University of Connecticut’s Center for Genome Innovation. Specifically, cDNA library preparation was performed using the Illumina TruSeq Stranded Total RNA Library Sample Prep Kit (RS-122-2201) with RiboZero for ribosomal RNA depletion. Sequencing was performed on the Illumina Novaseq 6000 platform. Reads were mapped to the mm10 genome (UCSC genome browser) using Hisat2^54^. Isoform and gene expression was determined through IsoEM2, to produce expression values in TPM^55^; differential gene expression was then determined using IsoDE2, as previously described^40^. Minor intron retention and ORF analyses were performed as described in Baumgartner et al. 2018, while alternative splicing around the minor intron was calculated and assessed as described in Olthof et al. 2021^40,44^. DAVID was employed for functional enrichment analysis of gene sets, with a significance cut-off of Benjamini-Hochberg adjusted *P*-value<0.05^62^

### Immunofluorescence

For IF, 16 µm coronal cryosections (E10.5-E12.5) were used as described previously^63^. Primary antibodies were diluted to to 1:200 (rabbit anti-NeuN, Abcam, ab17787) or 1:300 (rabbit anti-RFP, Rockland, 600-401-379; chicken anti-RFP, Rockland, 600-901-379; rabbit anti-PH3, Bethyl Laboratories, IHC-00061; rabbit anti-cleaved caspase 3, Cell Signaling Technology, 9665S; mouse anti-KI67, BD Biosciences, 556003).

### Whole-mount *in situ* hybridization (WISH)

WISH analysis was carried out as described previously^61,64^. E12.5 embryos were halved with tungsten needles (Fine Science Tools, 10130-10) after rehydration. Probes were generated using the primers listed in **Supplementary Data Table 8**.

### ImageJ

ImageJ to measure the tdTomato+ neuroectodermal length of ventricle to normalize embryonic PH3 quantification^65^ (**Fig. 1H**) and body lengths of P30 control and mutants across both sexes to evaluate stature^60^(**Supplementary Data 6A**,**B**).

### Neurolucida 3D hypothalamic reconstruction

Alternating 50µm coronal sections were imaged, and the contours of the tdTomato+ hypothalamic footprint and surface of the brain were traced to generate 3D reconstructions. Volume and surface area analysis were performed using StereoInvestigator and Neurolucida Explorer software (MBF Bioscience).

### Reconstruction

Serial images of 25µm thick coronal sections were first scaled, then aligned, and then the whole brain and *in situ* positivity was traced to generate 3D reconstructions with Reconstruct software^59^.

### Nissl staining

Postnatal day (P) 30 brains were cryosectioned (50 µm) and collected on 1% gelatin and chrome alum-coated glass slides. Slides were rinsed in distilled water for 1 min, followed by a sequence of ethanol (EtOH) dehydration steps for 2 min each (50% EtOH, 70% EtOH, 95% EtOH, 100% EtOH, 100% EtOH, 100% EtOH). This was followed by two 2-min toluene washes, and then by a sequence of dehydration steps in ethanol, each lasting 2 min (100% EtOH, 100% EtOH, 100% EtOH, 100% EtOH, 95% EtOH, 70% EtOH, 50% EtOH). Slides were rinsed in distilled water for 2 min, followed by 3 min in Cresyl Violet (1% cresyl violet+0.25% glacial acetic acid) and another 2 min water rinse. The first dehydration series was repeated, followed by three toluene washes (2 min, 1 min and 1 min). Slides were mounted with Permount mounting medium (Fisher Scientific, SP15-100) and allowed to harden in the fume hood overnight before being imaged and stitched with Keyence BZ-X710 at 4X.

### scRNAseq of juvenile hypothalamus

tdTomato+ *Nkx2*.*1*-*Cre* lineage was microsdissected from control (n=3) and mutant (n=3) juvenile mice for scRNAseq (P30; *N*=2, with each *N* containing 3 pooled hypothalami) and tissue was prepared for single cell dissociation as previously described^27^. Viability of single-cell suspensions was assessed with the BioRad TC20 Automated Cell Counter. Cells were then processed through the 10X Genomics Chromium Controller using the v3 chemistry kit. Specifically, post-cell capture and lysis, complementary DNA was synthesized and amplified (through 14 cycles) using the 10X Genomics protocol, after which a sequencing library was constructed for each sample. Libraries were sequenced on the Illumina NextSeq 550 platform using the v2.5 high-output sequencing kit. Sequencing data was first processed using the Cell Ranger pipeline (10X Genomics, Cellranger count v3.0.2) with the default parameters. Cells with >40% mitochondrial mitochondrial reads or less than 500 unique molecular identifiers (UMIs) were filtered. Further rounds of unsupervised and supervised clustering as well as data visualization were done using Seurat^56,58,66,67^.

### Liquid food consumption

Liquid food consumption was performed on singly housed control and mutant mice in a static mouse shoe box cage with sawdust bedding, enrichment (cotton nestlet, hut), water and liquid rodent food in a 50mL food dispenser (food: Bioserv. #9019; dispenser: F1268SP). Liquid food was prepared and stored according to manual. Mice were weaned from mothers at P21 and were provided 1-week to acclimate to liquid food diet before consumption was measured.

### Pair-feeding

Pair-feeding experiments were performed on age-matched control and mutant mice on singly housed animals using liquid food repeated through 11 weeks of age. After acclimation to liquid food diet post-weaning, *ad libitum* liquid food was provided to a control mouse. The amount of food consumed by the control mouse was measured over the course of two days and averaged. The average amount of food was then provided to the mutant mouse for two days, while the amount of food the control mouse ate was being measured and averaged.

### Metabolic Analyses by National Mouse Metabolic Phenotyping Center (UMass MMPC)

Assessment of whole-body fat, lean and water mass using was non-invasively measured in awake mice using 1H-MRS (Echo Medical System). Metabolic cages (TSE Systems) were used awake mice to simultaneously measure energy expenditure, physical activity, and indirect calorimetry. Here, the metabolic cages non-invasively measure O_2_ consumption and CO_2_ production in individual mice and calculates the respiratory exchange ratio to reflect energy expenditure. The metabolic cages will also be used for the quantitative measurement of horizontal and vertical movement (XYZ-axis) as an index of physical activity over 3 days. Assays for hormones (growth hormone, adrenocorticotropic hormone, prolactin, thyroid stimulating hormone, thyrotropin releasing hormone, and thyroxine) accessible online on National Mouse Metabolic Phenotyping Center (Experiment 22463).

### Feeding Experimentation device 3 (FED3)

*Free-feeding*. In the free-feeding mode (**Fig. 7A**) where upon pellet (20 mg 5TUM, Test Diet) retrieval from the pellet well, a subsequent pellet was dispensed^50^. Retrieval was monitored by infra-red beam break sensor across the pellet well. Latency to retrieval and time stamp were logged on an internal microSD. An experimental cohort of 16 female mice (8 control, 8 mutant) were removed from their group housing and singly house in static rat cages for a habituation period of 3-5 days on a 12/12 on/off light cycle. Mouse weight was monitored daily, and habitation continue until at least 120 pellets were retrieved in one 12-hour dark period, per correspondence with Dr. Alexxai V. Kravitz. Mutant mice occasionally did not learn to retrieve pellets from FED3. These mice were paired with control mice to habituate to the free-feeding paradigm. FED3 devices were checked for functionality, but mice were otherwise undisturbed. Free feeding was recorded for 24 hours.

*FED3 Data analysis*. FED3 data was downloaded as comma delineated CSV files and trimmed to exactly 24 hours. Fed3Viz was used to generate chronograms and bin pellet retrieval by 6-hour light cycle quarters. Also, meal size was binned by the number of pellets retrieved within a one-minute period. Binning data was downloaded from Fed3Viz and processed using GraphPad Prism.

### Imaging and quantification

P30 *in situ* and Nissl -processed slides were imaged and stitched with Keyence BZ-X710 at 4X. Whole mount *in situs* and P30 tdTomato+ images were taken using Olympus SZX10 Stereomicroscope. Embryonic immunofluorescence, *in situ*, and TUNEL-processed slides were imaged using a Leica SP2 confocal microscope, utilizing consistent laser intensity, excitation-emission windows, and gain and offset settings across control, control and mutant sections. For each channel, confocal imaging settings were optimized for fluorescence in the control section for each processed slide. Further processing was performed on IMARIS v8.3.1 (Bitplane) and Adobe Photoshop CS4 (Adobe Systems). As with the confocal settings, image processing was identical among images of control and mutant sections from the same slide. Manual quantification was performed in IMARIS as described in previously^40^.

### Statistical Tests

All statistical tests, n-values and comparisons made are presented in **Supplementary Data Table 9**.

## Data and code availability

The bulk RNA-sequencing and single cell RNA-sequencing data discussed in this publication have been deposited in NCBI’s Gene Expression Omnibus (GEO) and are accessible through GEO Series accession number GSE214104. Mouse phenotyping, performed by the National Mouse Metabolic Phenotyping Center at the University of Massachusetts, is available online experiment 22463. Any additional information required to reanalyze the data reported in this work paper is available from the lead contact upon request.

## Acknowledgments

The authors would like to thank Dr. Bo Reese from the Center of Genome Innovation at the University of Connecticut for performing the RNA sequencing. Additionally, we would like to acknowledge Dr. Ion Mandoiu from the Computer Science Engineering Department of the University of Connecticut for creating and maintaining the necessary infrastructure to perform bioinformatics analysis. Finally, we thank Dr. Linnaea Ostroff for her insightful comments on the manuscript. This work was supported by NIH R01NS102538 to RNK.

