## Supplemental Data for "Integrity of the minor spliceosome in the developing mouse hypothalamus determines neuronal subtype composition regulating energy balance"

**Supplementary figures and tables.**

**Supplementary Data Table 1 | Expression of select genes in control and mutant E12.5 ventral diencephalon.**

| | Gene | Control (TPM) | Mutant (TPM) |
|---|---|---|---|
| Genetics | *Rnu11* | 742.2629824 | 457.5505701 |
| | *Nkx2-1* | 64.89170326 | 75.02268457 |
| Expressed VD | *Rax* | 4.505701465 | 3.583683628 |
| | *Six3* | 11.66945909 | 10.9949361 |
| | *Nr5a1* | 6.300193998 | 5.746568103 |
| Non-Expressed VD | *Emx1* | 0.023871212 | 0.046463588 |
| | *Foxg1* | 0.20874984 | 0.095286747 |
| | *Ikzf1* | 0.090184004 | 0.493047213 |
| Downregulated VD markers | *Pomc* | 7.387132887 | 2.943294501 |
| | *Foxb1* | 7.648602963 | 4.666412787 |
| | *Neurog1* | 3.468572989 | 2.46262375 |
| Upregulated enriching for GOTERM: Synapse | *Actb* | 2239.475463 | 8746.669868 |
| | *Adgrl1* | 18.62854659 | 26.15441847 |
| | *Adra2a* | 0.936744999 | 1.617921444 |
| | *Apoe* | 8.624346953 | 33.43051366 |
| | *Asic1* | 5.13754717 | 7.923254186 |
| | *Asic2* | 2.192230866 | 4.849153749 |
| | *Atp2b2* | 4.887613123 | 8.106955454 |
| | *Cacng2* | 2.330625778 | 4.341998915 |
| | *Cartpt* | 2.320702705 | 6.907203764 |
| | *Cbarp* | 28.14850418 | 38.7121901 |
| | *Chrna5* | 0.452290562 | 1.034213706 |
| | *C1qa* | 1.068036195 | 5.614782113 |
| | *C1qb* | 2.249455654 | 10.35218554 |
| | *C1qc* | 1.591747236 | 7.273484322 |
| | *Gabra4* | 0.765237479 | 1.855664036 |
| | *Glra2* | 3.097048172 | 5.696952195 |
| | *Gria3* | 2.437320181 | 6.252909103 |
| | *Grik1* | 3.832202441 | 8.319866168 |
| | *Grin2d* | 1.154686322 | 2.263658147 |
| | *Lrrc4c* | 6.328302429 | 12.47818043 |
| | *Mctp1* | 0.685739471 | 1.513692005 |
| | *Mme* | 0.45418126 | 1.185861431 |
| | *Ppfia4* | 1.656294851 | 2.436828212 |
| | *Prrt1* | 4.888497153 | 6.748339354 |
| | *Slc4a10* | 1.092039243 | 2.407382908 |
| | *Snph* | 4.786651588 | 7.40183957 |
| | *Sv2b* | 1.358822281 | 3.128929251 |
| | *Syt7* | 9.645691158 | 17.9761883 |
| | *Tpd52* | 5.846885831 | 12.45126025 |
| | *Unc13c* | 0.679804096 | 2.052966382 |

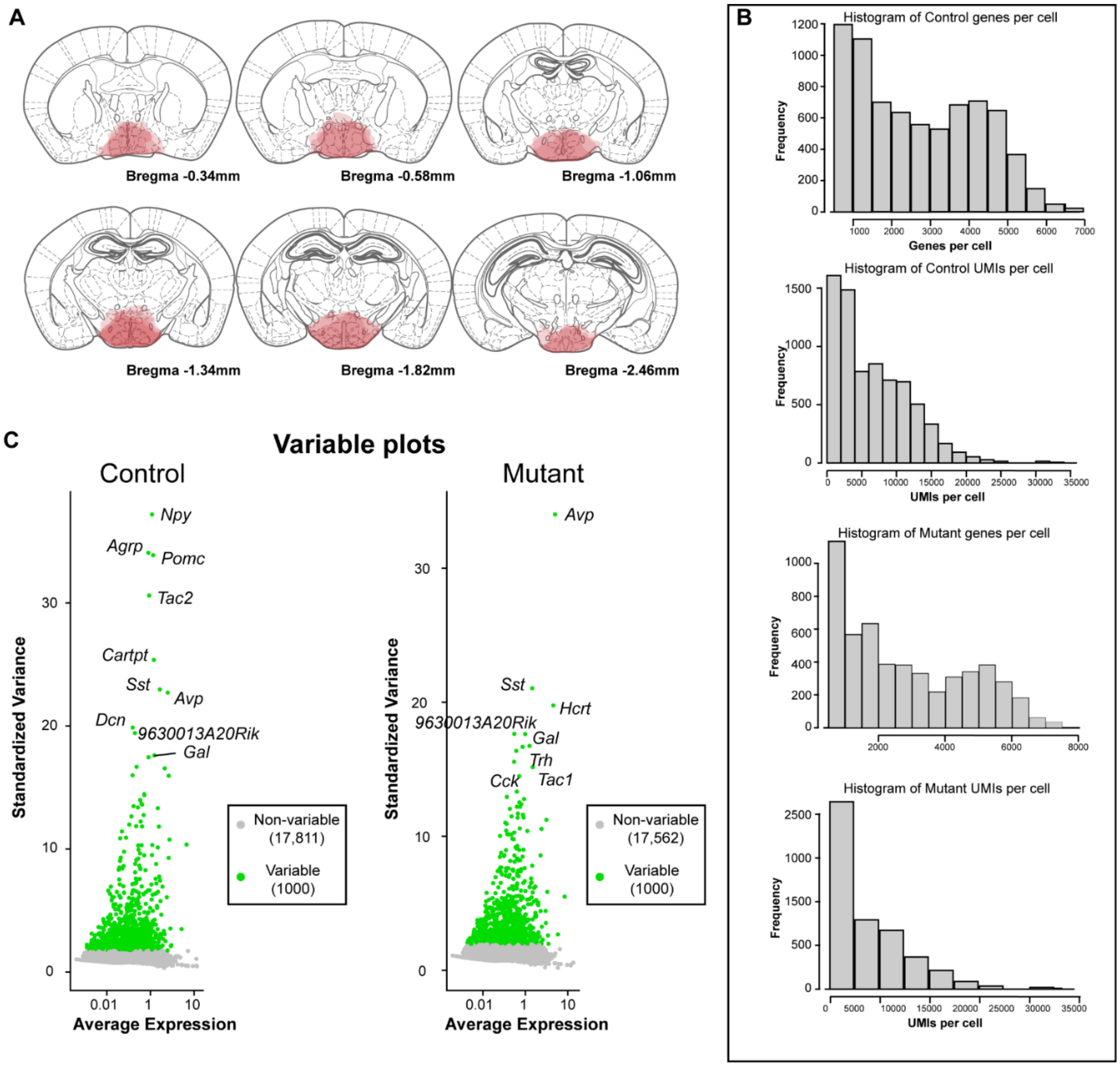

**Supplementary Data 2 | *Nkx2.1*-derived hypothalamic scRNAseq methods and descriptive data.**

(**a**) Schematics of hypothalamic tissue punches with Bregma coordinates used for scRNAseq isolation and analysis. (**b**) Histogram of the number of genes per cell and unique molecular identifiers (UMIs) per cell in control (top) and mutant (bottom) data sets. (**c**) Top 10 variable genes on control (left) and mutant (right). Non-variable genes=grey; Variable genes=green.

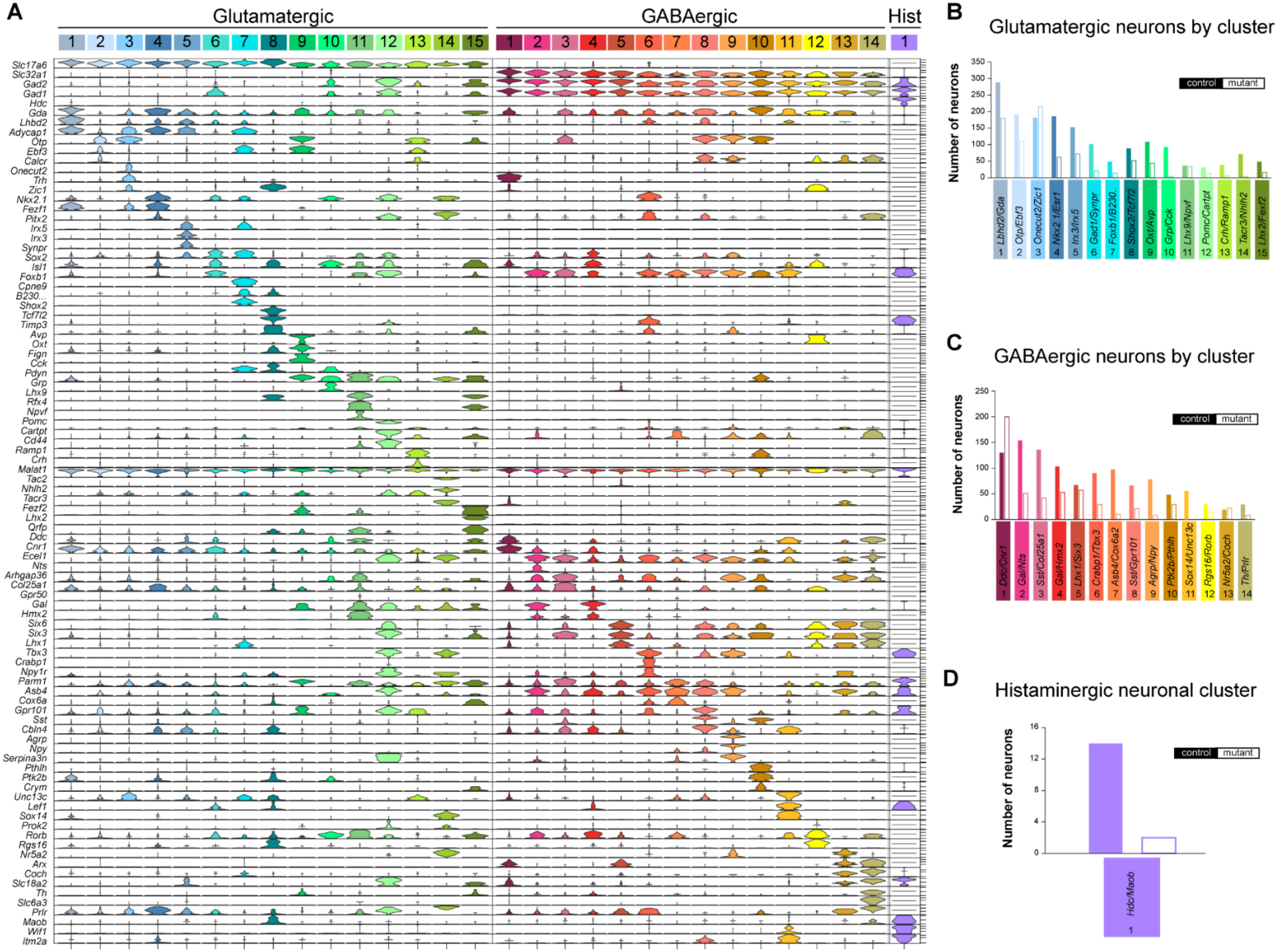

**Supplementary Data 3 | Violin plot and numeric breakdown by genotype of identified neuronal clusters by scRNAseq neuronal clusters**. (**a**) Violinplots of the top 3 genes expressed per cluster by Glutamatergic, GABAergic, and Histaminergic neuronal cell types identified. (**b-d**) Bar charts of the number of control (filled bar) and mutant (empty bar) neurons per cluster for Glutamatergic (**b**), GABAergic (**c**), and histaminergic (**d**) neurons.

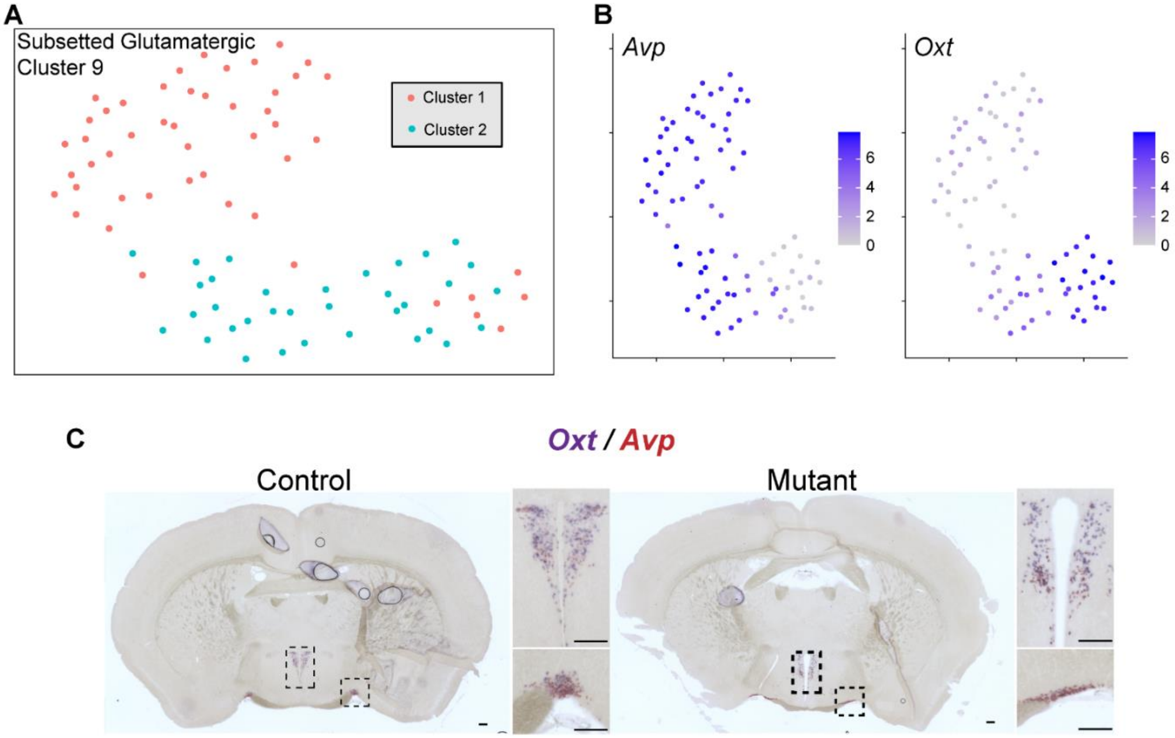

**Supplementary Data 4 | Sub-setted Glutamatergic Cluster 9 reveals *Oxt*⁺/*Avp*⁻ and *Avp*⁺/*Oxt*⁻ sub-populations.**

(**a**) UMAP of sub-setted Clutamatergic Cluster 9, color coded by cluster. (**b**) Feature plots of *Avp* (left) and *Oxt* (right). (**c**) Double *in situ* hybridization for *Oxt* (purple) and *Avp* (red) on control (left) and mutant (right) P30 brains. Insets of paraventricular (top) and suprachiasmatic (bottom) nuclei. Scale bar 200µm.

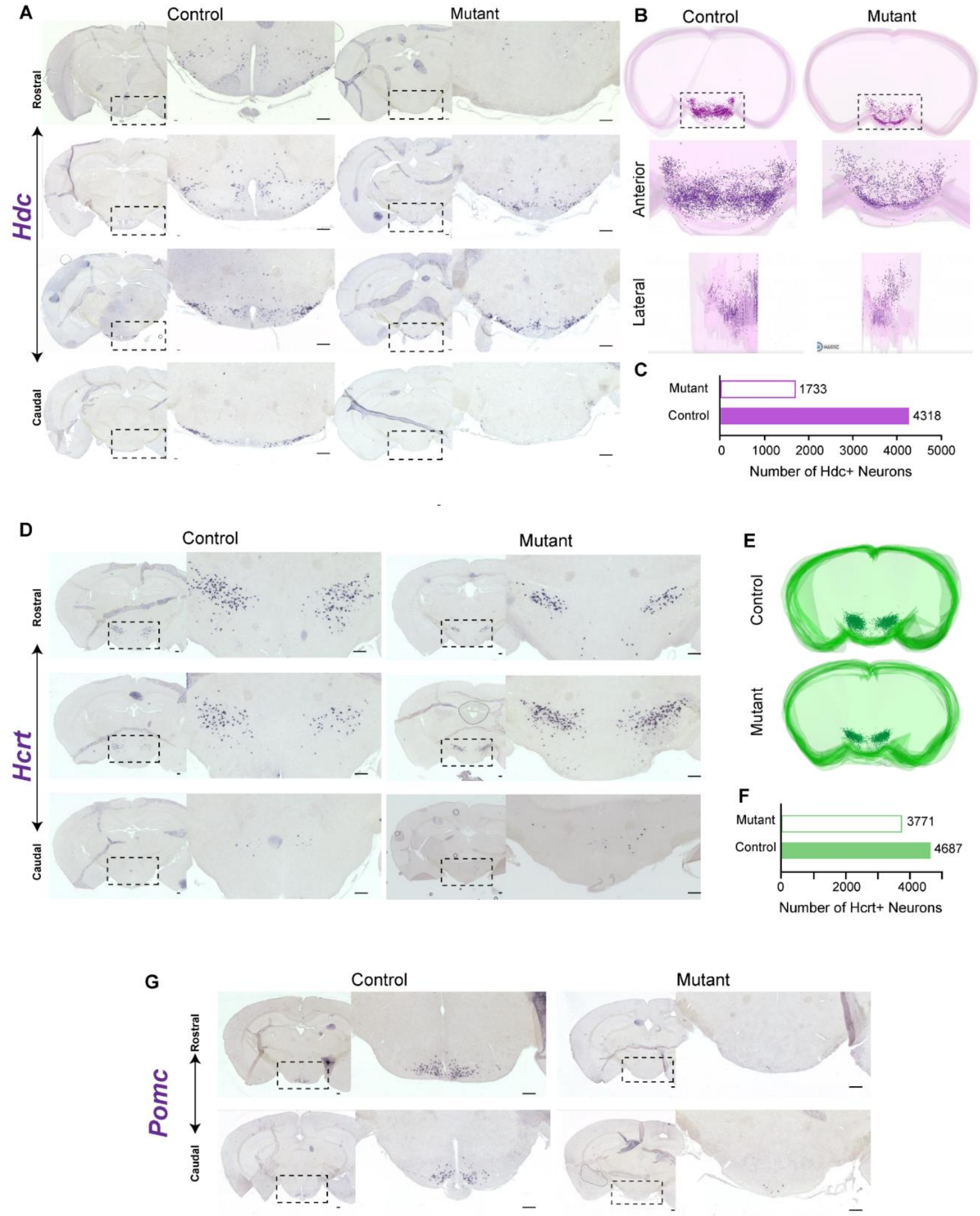

**Supplementary Data 5 | Validations of reduced histaminergic and *Pomc*+ neurons, comparable *Hcrt* neurons in the *Rnu11* cKO mouse.**

(**a**) Representative images from rostral (top) to caudal (bottom) of *in situ* hybridization (ISH) on coronal sections on control (left) and mutant (right) brain for *Hdc*. (**b**) 3D rendering of *Hdc* ISH in P30 juvenile control and mutant. Zoomed inset of 3D reconstruction from anterior and lateral views. *Hdc*$^{+}$ neurons = purple dots. (**c**) Quantification of *Hdc* neurons in control and mutant P30 hypothalamii (n=1, serial sections across full hypothalamus). (**d**) Representative images from rostral (top) to caudal (bottom) of ISH on coronal sections on control (left) and mutant (right) brain for *Hcrt*. (**e**) 3D rendering of *Hcrt* ISH in P30 juvenile control and mutant. Zoomed inset of 3D reconstruction from anterior and lateral views. *Hdc*$^{+}$ neurons = green dots. (**f**) Quantification of *Hcrt* neurons in control and mutant P30 hypothalamii (n=1, serial sections across full hypothalamus). (**g**) Representative images from rostral (top) to caudal (bottom) of ISH on coronal sections on control (left) and mutant (right) brain for *Pomc*. Scale bar 200µm.

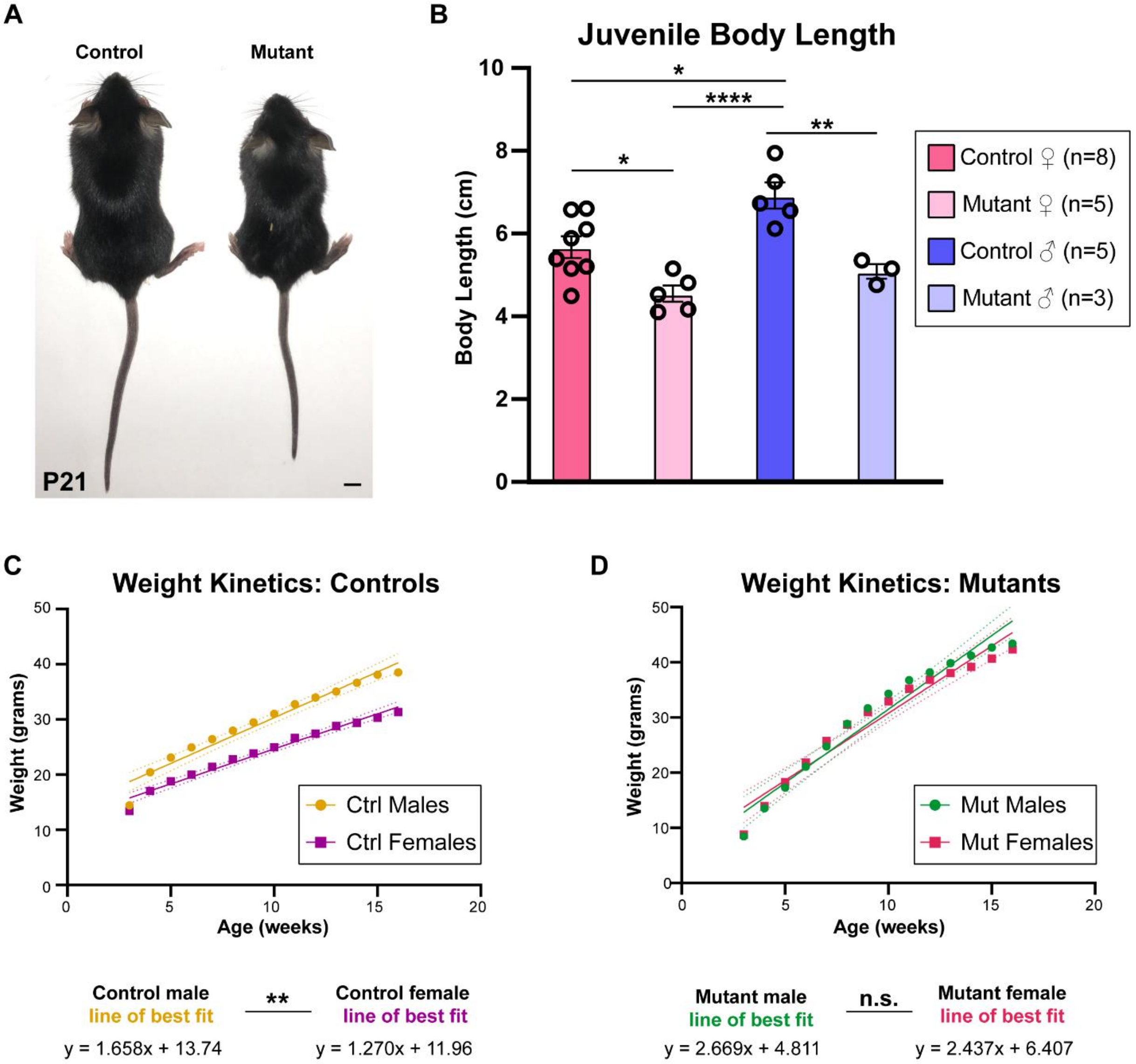

**Supplementary Data 6 | Postnatal phenotype of short stature at weaning age and weight kinetics analysis.**

(**a**) Representative image of P21 control (left) and mutant (right) females. Scale bar 0.5 cm. (**b**) Bar chart of juvenile body lengths of control females (dark pink, n=8), mutant females (light pink, n=5), control males (violet, n=5), and mutant males (light violet, n=3). (**c**) Weight kinetics of control male (orange circles) and control female (purple boxes). Lines of best fit (solid line) and the 95% confidence interval (dotted lines) per condition. (**d**) Weight kinetics of mutant male (green circles) and mutant female (magenta boxes). Lines of best fit (solid line) and the 95% confidence interval (dotted lines) per condition. Data (**a,b**) are presented as mean±s.e.m. Statistical significance determined by one-way ANOVA followed by post-hoc Tukey test. ****=*P*<0.0001, **=*P*<0.01, *=*P*<0.05. Simple linear regression to test for significantly different slopes (See also **Supplementary Data Table 9**).

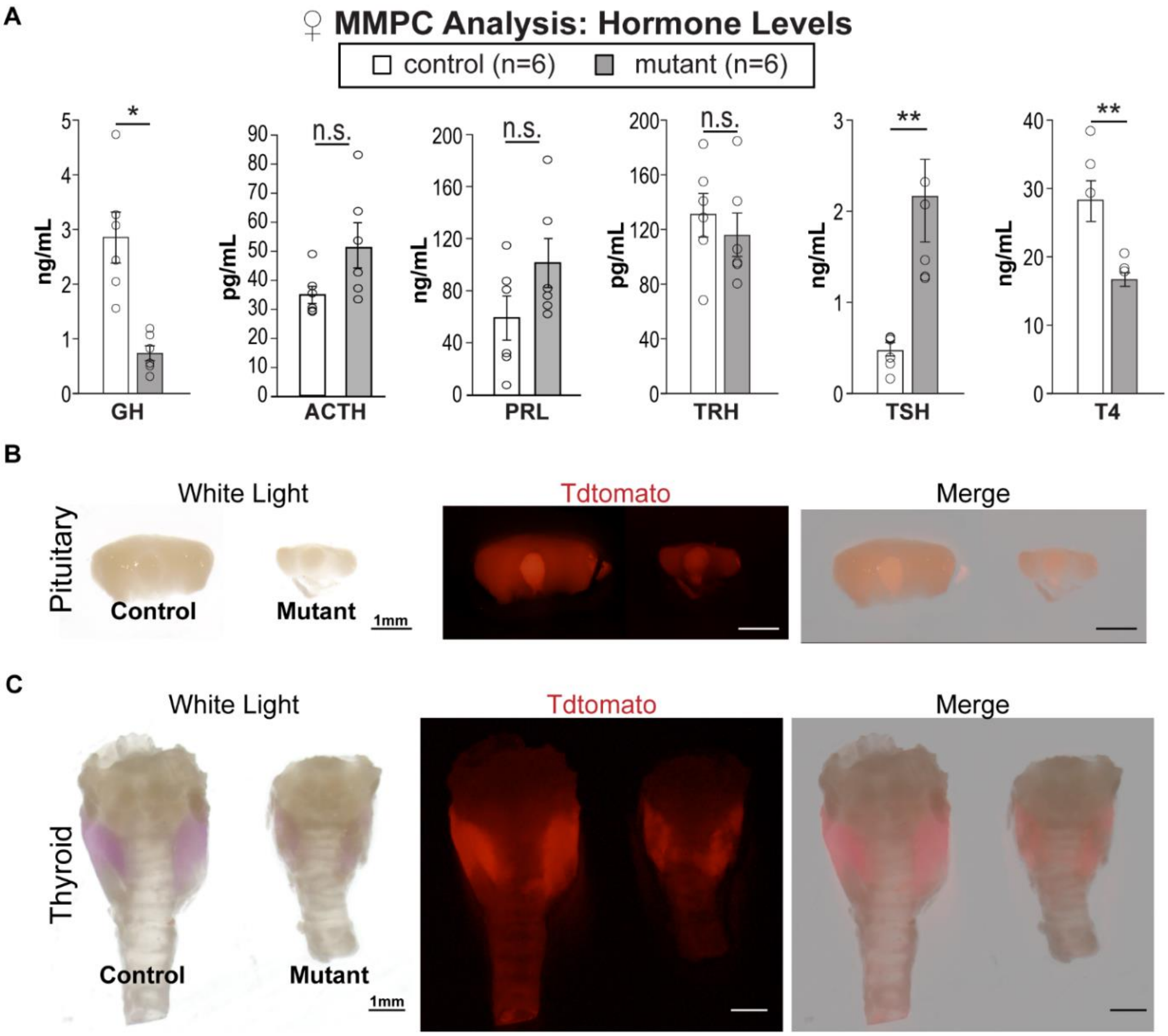

**Supplementary Data 7 | Hormonal defects in *Rnu11*-null *Nkx2.1*-Cre+ mice.**

(**a**) Bar charts of circulating hormone analysis for pituitary hormones (growth hormone (GH), adrenocorticotropic hormone (ACTH), prolactin (PRL) and thyroid stimulating hormone (TSH)), hypothalamic hormone (thyrotropin releasing hormone (TRH)), and thyroxine (T4), for controls (white bars) and mutants (grey bars). (**b**) White light (left), tdTomato (middle) and merged images of P30 pituitaries from control (left) and mutant (right). (**c**) White light (left), tdTomato (middle) and merged images of P30 thyroids from control (left) and mutant (right). Scale bar 1mm. Scale bar 1mm. Data represented as mean, with error bars ±SEM. Significance was determined by two-tailed student's t-tests. n.s.= not significant, *=$P$<0.05, **=$P$<0.01, ***=$P$<0.001, ****=$P$<0.0001 (See also **Supplementary Data Table 9**).

**Supplementary Data Table 8 | Primers used for genotyping and *in situ* hybridization.**

| Primer Name | Forward | Reverse | Referenced |
|---|---|---|---|
| *Rnu11* Genotyping | ATAGTGCATGAATAGAGCACCCTCCC | AGGCTGCTACAGGATGACTCTGTCTTCT | Methods |
| Cre Genotyping | TATCCAGCAACATTTGGGCCAGCT | AACATTCTCCCACCGTCAGTACGTGA | Methods |
| tdTomato WT Genotyping | AAGGGAGCTGCAGTGGAGTA | CCGAAAATCTGTGGGAAGTC | Methods |
| tdTomato MUT Genotyping | GGCATTAAAGCAGCGTATCC | CTGTTCCTGTACGGCATGG | Methods |
| *Rnu11* | AAAGGGCTTCTGTCGTGAGTGGC | CCGGGACCAACGATCACCAG | Figures 1; 2 |
| *Nkx2.1* | CTTCTTCTTCTCCTCCTCCTCC | TGAAAAAGTGAGGGACTAGGGA | Figure 2 |
| *Pomc* | CGACGGAAGAGAAAAGAGGTTA | CTTGGAATGAGAAGACCCCTG | Figures 2,5; Suppl. Data 5 |
| *Lef1* | AGAGAACACCCTGATGAAGGAA | CTTCCTCTTCTTCTTCTTGCCA | Figure 2 |
| *Lhx6* | TCAACAACAGCCTCCTACTTCA | TCTCTCCATGAAACTCTCGTCA | Figure 2 |
| *Shh* | ATGCTGCTGCTGCTGGCCAGATGT | GGGCCCCGAGTCGTTGTGCGGCGC | Figure 2 |
| *Foxb1* | GCCTATCCTCTCCCTAACCAGT | TTTCAGGACCTAAGAAGATCCG | Figure 2 |
| *Irx5* | AGAAAGAGAACAAGATGACGTGG | CGAAGGAGCCATAGTTCGTGTA | Figure 2 |
| *Hcrt* | TTCCTTCTACAAAGGTTCCCTG | CAGGACAAGGATAGAAGATGGG | Suppl. Data 5 |
| *Hdc* | TCAGTGGAGAAGGCTGGC | TCATCTCCTCCCTCGCTG | Suppl. Data 5 |
| *Npy* | CTTGAAGACCCTTCCATGTGGTGA | TGGGACAGGCAGACTGGTTTCAG | Figure 5 |
| *Oxt* | CTGGATATGCGCAAGTGTCTC | AAAGGTATTCCCAGAAAGTGGG | Suppl. Data 4 |
| *Avp* | CCGAGTGCCACGACGGTTT | TTCCATGCTGTAGGGGCGAG | Suppl. Data 4 |

**Supplementary Data Table 9 | Results of all statistical tests.**

| Figure Panel | Analysis | Comparison | N-Value | Timepoint | Control | Mutant | Statistical test | P-Value | * |
|---|---|---|---|---|---|---|---|---|---|
| 1.e. | Minor intron retention (median mis-splicing Index) | control vs mutant | 409 minor introns | E12.5 | 0.106171494 | 0.162993499 | Mann-Whitney U-test | P<0.00001 | **** |
| 1.h. | PH3/DAPI per tdt+ ventricular length | control vs mutant | 3 | E10.5 | 0.0882±0.0181 | 0.1352±0.0366 | Student's two-tailed T-test | P =0.3938 | n.s. |
| 1.h. | PH3/DAPI per tdt+ ventricular length | control vs mutant | 4 | E11.5 | 0.1586±0.0305 | 0.2515±0.0239 | Student's two-tailed T-test | 0.0743 | n.s. |
| 1.h. | PH3/DAPI per tdt+ ventricular length | control vs mutant | 3 | E12.5 | 0.0611±0.0109 | 0.2562±0.0619 | Student's two-tailed T-test | 0.0361 | * |
| 1.j. | CC3+ cells in 100 µm column | control vs mutant | 3 | E10.5 | 1.6667±1.6667 | 6±5.0332 | Student's two-tailed T-test | 0.9905 | n.s. |
| 1.j. | CC3+ cells in 100 µm column | control vs mutant | 4 | E11.5 | 0.2500±0.2500 | 50±7.8422 | Student's two-tailed T-test | <0.0001 | **** |
| 1.j. | CC3+ cells in 100 µm column | control vs mutant | 3 | E12.5 | 0±0 | 60.3333±6.4377 | Student's two-tailed T-test | <0.0001 | **** |
| 1.j. | TUNEL+ cells in 100 µm column | control vs mutant | 3 | E10.5 | 3±1.528 | 25±6.807 | Student's two-tailed T-test | 0.9241 | n.s. |
| 1.j. | TUNEL+ cells in 100 µm column | control vs mutant | 4 | E11.5 | 2.5±0.866 | 69±10.512 | Student's two-tailed T-test | 0.0448 | * |
| 1.j. | TUNEL+ cells in 100 µm column | control vs mutant | 3 | E12.5 | 0±0 | 154.3333±39.725 | Student's two-tailed T-test | 0.0001 | *** |

| | | | | | | | | | |
|---|---|---|---|---|---|---|---|---|---|
| 2.e. | NeuN/Ki67 ratio | control vs mutant | 3 | E11.5 | 0.3352 ± 0.05053 | 0.7572 ± 0.2653 | Student's two-tailed T-test | P =0.1932 | n.s. |
| 2.e. | NeuN/Ki67 ratio | control vs mutant | 4 | E12.5 | 0.5781 ± 0.1855 | 1.386 ± 0.2341 | Student's two-tailed T-test | P =0.0353 | * |
| 3.b'. | Normalized Tdt+volume (AU) | control vs mutant | 3 | juvenile | 1 ± 0.0278 | 0.7680 ± 0.02712 | Student's two-tailed T-test | P = 0.0040 | ** |
| 6. g. | Weight Kinetics Females (grams) | control vs mutant | control=9; mutant=11 | 3 weeks | 13.4444 ± 0.5411 | 8.7990 ± 0.4523 | Student's two-tailed T-test | P = <0.0001 | **** |
| | | control vs mutant | | 4 weeks | 17.0833 ± 0.4692 | 13.9909 ± 0.7046 | Student's two-tailed T-test | P = 0.0027 | ** |
| | | control vs mutant | | 5 weeks | 18.8 ± 0.5875 | 18.3136 ± 1.1328 | Student's two-tailed T-test | P = 0.7257 | n.s. |
| | | control vs mutant | | 6 weeks | 20.0122 ± 0.7955 | 21.8527 ± 1.5307 | Student's two-tailed T-test | P = 0.3317 | n.s. |
| | | control vs mutant | | 7 weeks | 21.4444 ±1.0028 | 25.7663 ± 1.5194 | Student's two-tailed T-test | P = 0.0366 | * |
| | | control vs mutant | | 8 weeks | 22.8178 ± 1.2298 | 28.71 ± 1.3228 | Student's two-tailed T-test | P = 0.0049 | ** |
| | | control vs mutant | | 9 weeks | 23.84 ± 1.2860 | 30.9791 ± 1.3427 | Student's two-tailed T-test | P = 0.0014 | ** |
| | | control vs mutant | | 10 weeks | 24.99 ± 1.4658 | 32.9273 ± 1.4331 | Student's two-tailed T-test | P = 0.0012 | ** |
| | | control vs mutant | | 11 weeks | 26.6533 ± 1.7027 | 35.2364 ± 1.5344 | Student's two-tailed T-test | P = 0.0015 | ** |
| | | control vs mutant | | 12 weeks | 27.4256 ± 1.7671 | 36.8709 ± 1.6411 | Student's two-tailed T-test | P = 0.0010 | ** |

| | | | | | | | | | |
|---|---|---|---|---|---|---|---|---|---|
| | | control vs mutant | | 13 weeks | 28.8056 ± 1.9357 | 38.0782 ± 1.6887 | Student's two-tailed T-test | P = 0.0019 | ** |
| | | control vs mutant | | 14 weeks | 29.3789 ± 2.1564 | 39.17 ± 1.6181 | Student's two-tailed T-test | P = 0.0016 | ** |
| | | control vs mutant | | 15 weeks | 30.3256 ± 2.3765 | 40.7064 ± 1.8951 | Student's two-tailed T-test | P = 0.0028 | ** |
| | | control vs mutant | | 16 weeks | 31.3456 ± 2.6118 | 42.3745 ± 1.8796 | Student's two-tailed T-test | P = 0.0025 | ** |
| 6. h. | Weight Kinetics Males (grams) | control vs mutant | control=7; mutant=8 | 3 weeks | 14.4229 ± 0.8785 | 8.475 ± 0.9514 | Student's two-tailed T-test | P = 0.0005 | *** |
| | | control vs mutant | | 4 weeks | 20.4571 ± 0.5323 | 13.5625 ± 1.0188 | Student's two-tailed T-test | P = <0.0001 | **** |
| | | control vs mutant | | 5 weeks | 23.0871 ± 0.5634 | 17.3325 ± 1.2228 | Student's two-tailed T-test | P = 0.0013 | ** |
| | | control vs mutant | | 6 weeks | 24.9214 ± 0.5155 | 21.1188 ± 1.4563 | Student's two-tailed T-test | P = 0.0370 | * |
| | | control vs mutant | | 7 weeks | 26.4271 ± 0.6649 | 24.7913 ± 1.7902 | Student's two-tailed T-test | P = 0.4326 | n.s. |
| | | control vs mutant | | 8 weeks | 27.9757 ± 0.8584 | 28.8488 ± 1.8843 | Student's two-tailed T-test | P = 0.6948 | n.s. |
| | | control vs mutant | | 9 weeks | 29.49 ± 0.9783 | 31.6988 ± 1.7991 | Student's two-tailed T-test | P = 0.3200 | n.s. |
| | | control vs mutant | | 10 weeks | 31.0071 ± 1.1265 | 34.325 ± 1.7142 | Student's two-tailed T-test | P = 0.1414 | n.s. |
| | | control vs mutant | | 11 weeks | 32.7714 ± 1.3676 | 36.7875 ± 1.5352 | Student's two-tailed T-test | P = 0.0760 | n.s. |

| | | | | | | | | | |
|---|---|---|---|---|---|---|---|---|---|
| | | control vs mutant | | 12 weeks | 33.9786 ± 1.4069 | 38.1688 ± 1.5459 | Student's two-tailed T-test | P = 0.0690 | n.s. |
| | | control vs mutant | | 13 weeks | 35.0571 ± 1.5824 | 39.8625 ± 1.4542 | Student's two-tailed T-test | P = 0.0433 | * |
| | | control vs mutant | | 14 weeks | 36.6571 ± 1.5085 | 41.2425 ± 1.3270 | Student's two-tailed T-test | P = 0.0392 | * |
| | | control vs mutant | | 15 weeks | 38.0914 ± 1.5374 | 42.6875 ± 1.2802 | Student's two-tailed T-test | P = 0.0375 | * |
| | | control vs mutant | | 16 weeks | 38.5371 ± 1.7410 | 43.4163 ± 1.3263 | Student's two-tailed T-test | P = 0.0415 | * |
| 6. i. | Physical activity (day) (counts) | control vs mutant | 6 | 13 weeks | 3207.6667 ± 278.9115 | 1210.8333 ± 76.0918 | Student's two-tailed T-test | P = <0.0001 | **** |
| | Physical activity (night) (counts) | control vs mutant | 6 | 13 weeks | 8178.6667 ± 608.8510 | 1624.1667 ± 98.2892 | Student's two-tailed T-test | P = <0.0001 | **** |
| | Physical activity (24hr) (counts) | control vs mutant | 6 | 13 weeks | 5692.8333 ± 432.2789 | 1417.6667 ± 80.6109 | Student's two-tailed T-test | P = <0.0001 | **** |
| 6. j. | Oxygen consumption (day) | control vs mutant | 6 | 13 weeks | 3725.1667 ± 86.884 | 2885.3333 ± 75.964 | Student's two-tailed T-test | P = <0.0001 | **** |
| | Oxygen consumption (night) | control vs mutant | 6 | 13 weeks | 4286.5 ± 127.1162 | 2857.5 ± 81.4312 | Student's two-tailed T-test | P = <0.0001 | **** |
| | Oxygen consumption (24hr) | control vs mutant | 6 | 13 weeks | 4006 ± 103.340 | 2871.3333 ± 77.381 | Student's two-tailed T-test | P = <0.0001 | **** |
| 6. k. | Energy expenditure (day) | control vs mutant | 6 | 13 weeks | 18.4 ± 0.463 | 14.133 ± 0.415 | Student's two-tailed T-test | P = <0.0001 | **** |
| | Energy expenditure (night) | control vs mutant | 6 | 13 weeks | 21.3167 ± 0.665 | 14.1167 ± 0.442 | Student's two-tailed T-test | P = <0.0001 | **** |

| | | | | | | | | | |
|---|---|---|---|---|---|---|---|---|---|
| | Energy expenditure (24hr) | control vs mutant | 6 | 13 weeks | 19.8667 ± 0.556 | 14.1333 ± 0.426 | Student’s two-tailed T-test | P = <0.0001 | **** |
| 6. l. | RER (day) | control vs mutant | 6 | 13 weeks | 0.8992 ± 0.0126 | 0.8632 ± 0.0175 | Student’s two-tailed T-test | P = 0.1254 | n.s. |
| | RER (night) | control vs mutant | 6 | 13 weeks | 0.9277 ± 0.0143 | 0.8968 ± 0.0210 | Student’s two-tailed T-test | P = 0.2532 | n.s. |
| | RER (24hr) | control vs mutant | 6 | 13 weeks | 0.9137 ± 0.0129 | 0.88 ± 0.0185 | Student’s two-tailed T-test | P = 0.1657 | n.s. |
| 6. m. | Body weight (grams) | control vs mutant | 6 | 13 weeks | 23.6333 ± 0.8151 | 27.3167 ± 1.8452 | Student’s two-tailed T-test | P = 0.0978 | n.s. |
| | Fat mass weight (grams) | control vs mutant | 6 | 13 weeks | 3.4833 ± 0.3851 | 12.8667 ± 1.2505 | Student’s two-tailed T-test | P = <0.0001 | **** |
| | Lean mass weight (grams) | control vs mutant | 6 | 13 weeks | 18.9833 ± 0.6544 | 13.75 ± 0.8801 | Student’s two-tailed T-test | P = 0.0008 | *** |
| 6. n. | Body composition (fat to body mass) | control vs mutant | 6 | 13 weeks | 0.15 ± 0.0224 | 0.4672 ± 0.0225 | Student’s two-tailed T-test | P = <0.0001 | **** |
| | Body composition (lean to body mass) | control vs mutant | 6 | 13 weeks | 0.8028 ± 0.0123 | 0.5069 ± 0.0194 | Student’s two-tailed T-test | P = <0.0001 | **** |
| 6. p. | Liquid food consumption (mL) | control vs mutant | 6 | 4 weeks | 13.855 ± 0.3824 | 17.7633 ± 1.1735 | Student’s two-tailed T-test | P =0.0100 | * |
| | | control vs mutant | 6 | 5 weeks | 14.265 ± 0.6283 | 18.8067 ± 1.2579 | Student’s two-tailed T-test | P =0.0090 | ** |
| | | control vs mutant | 6 | 6 weeks | 15 ± 0.8318 | 19.0467 ± 0.4928 | Student’s two-tailed T-test | P =0.0019 | ** |
| | | control vs mutant | 6 | 7 weeks | 14.335 ± 1.0442 | 20.3333 ± 1.6363 | Student’s two-tailed T-test | P =0.0114 | * |

| | | | | | | | | | |
|---|---|---|---|---|---|---|---|---|---|
| 6. q. | Pair-feeding weight gain (grams) | control vs mutant | 3 | 5 weeks | 18.65 ± 0.3329 | 13.25 ± 1.0563 | Student's two-tailed T-test | P =0.0082 | ** |
| | | control vs mutant | 3 | 6 weeks | 20.8 ± 0.6007 | 15.9167 ± 1.3148 | Student's two-tailed T-test | P =0.0278 | * |
| | | control vs mutant | 3 | 7 weeks | 22.25 ± 1.2413 | 17.55 ± 1.4396 | Student's two-tailed T-test | P =0.0688 | n.s. |
| | | control vs mutant | 3 | 8 weeks | 23.0667 ± 1.4726 | 19.0167 ± 1.7742 | Student's two-tailed T-test | P =0.1538 | n.s. |
| | | control vs mutant | 3 | 9 weeks | 23.9833 ± 1.4990 | 21.05 ± 2.2567 | Student's two-tailed T-test | P =0.3398 | n.s. |
| | | control vs mutant | 3 | 10 weeks | 25.9667 ± 2.2556 | 22.5833 ± 2.5140 | Student's two-tailed T-test | P =0.3732 | n.s. |
| | | control vs mutant | 3 | 11 weeks | 26.55 ± 2.2605 | 23.9833 ± 2.0685 | Student's two-tailed T-test | P =0.4494 | n.s. |
| 7.b. | Pellets Retrieved Day | control vs mutant | 8 | Adult | 39.7770±7.0237 | 119.2553±18.6096 | Student's two-tailed T-test | 0.0013 | ** |
| 7.b. | Pellets Retrieved Night | control vs mutant | 8 | Adult | 144.4003±32.7480 | 169.3462±16.6475 | Student's two-tailed T-test | 0.2082 | n.s. |
| 7.c. | Average meal size histogram 1 pellet meal | control vs mutant | 8 | Adult | 38.8750±7.7769 | 59.0000±19.0066 | Student's two-tailed T-test | 0.3437 | n.s. |
| 7.c. | Average meal size histogram 2 pellet meal | control vs mutant | 8 | Adult | 18.1250±3.1816 | 34.1250±8.9611 | Student's two-tailed T-test | 0.1146 | n.s. |
| 7.c. | Average meal size histogram 3 pellet meal | control vs mutant | 8 | Adult | 7.8750±1.6084 | 19.8750±3.6422 | Student's two-tailed T-test | 0.0093 | * |
| 7.c. | Average meal size | control vs mutant | 8 | Adult | 6.1250±1.9588 | 12.6250±2.7963 | Student's two-tailed T-test | 0.0777 | n.s. |

| | | | | | | | | | |
|---|---|---|---|---|---|---|---|---|---|
| | histogram 4 pellet meal | | | | | | | | |
| 7.c. | Average meal size histogram 5 pellet meal | control vs mutant | 8 | Adult | 2.1250±0.9149 | 5.7500±3.4628 | Student's two-tailed T-test | 0.3287 | n.s. |
| 7.c. | Average meal size histogram 6 pellet meal | control vs mutant | 8 | Adult | 0.8750±0.6391 | 3.1250±2.0392 | Student's two-tailed T-test | 0.3102 | n.s. |
| 7.c. | Average meal size histogram 7 pellet meal | control vs mutant | 8 | Adult | 0.2500±0.1637 | 0.5000±0.2673 | Student's two-tailed T-test | 0.4384 | n.s. |
| 7.c. | Average meal size histogram 8 pellet meal | control vs mutant | 8 | Adult | 0.2500±0.2500 | 0.3750±0.2631 | Student's two-tailed T-test | 0.7356 | n.s. |
| 7.d. | Chronogram Q1 | control vs mutant | 8 | Adult | 17.6250±2.9212 | 51.1250±9.0661 | Student's two-tailed T-test | 0.0079 | ** |
| 7.d. | Chronogram Q2 | control vs mutant | 8 | Adult | 22.8750±4.5960 | 79.6250±11.2471 | Student's two-tailed T-test | 0.004 | *** |
| 7.d. | Chronogram Q3 | control vs mutant | 8 | Adult | 62.3750±8.0311 | 84.8750±9.1991 | Student's two-tailed T-test | 0.867 | n.s. |
| 7.d. | Chronogram Q4 | control vs mutant | 8 | Adult | 41.7500±6.1810 | 77.7500±8.4995 | Student's two-tailed T-test | 0.0041 | ** |
| 7.e. | Control consumption across quadrants | Q1 vs Q2 vs Q3 vs Q4 | 8 | Adult | see above for center value | | one way ANOVA, post-hoc Tukeys | Q1 v Q2 P= 0.3070; Q1 v Q3 P = 0.0011; Q1 v Q4 P = 0.0663; Q2 v Q3 P = 0.0044; Q2 v Q4 P = 0.3060; Q3 v Q4 P = 0.2786 | Q1 v Q2 P= n.s.; Q1 v Q3 P = **; Q1 v Q4 P =n.s.; Q2 v Q3 P = **; |

| | | | | | | | | | |
|---|---|---|---|---|---|---|---|---|---|
| | | | | | | | | | Q2 v Q4 P = n.s.; Q3 v Q4 P = n.s. |
| | Mutant consumption across quadrants | Q1 vs Q2 vs Q3 vs Q4 | 8 | Adult | | see above for center value | one way ANOVA, post-hoc Tukeys | Q1 v Q2 P=0.0022; Q1 v Q3 P = 0.0336; Q1 v Q4 P = 0.0219; Q2 v Q3 P = 0.9373; Q2 v Q4 P = 0.9962; Q3 v Q4 P = 0.7513 | Q1 v Q2 P= **; Q1 v Q3 P = *; Q1 v Q4 P = *; Q2 v Q3 P = n.s.; Q2 v Q4 P =n.s.; Q3 v Q4 P = n.s. |
| Suppl. data 6b | Juvenile Stature (cm) | control vs mutant including both sexes | females control=8; mutant=5 | ~P30 | 5.678 ± 0.263 | 4.5522 ± 0.197 | one way ANOVA, post-hoc Tukeys | control ♀ v mutant ♀ P= 0.0288; control ♀ v control ♂ P = 0.0147; control ♀ v mutant ♂ P = 0.5311; mutant ♀ v control ♂ P = <0.0001; mutant ♀ v mutant ♂ | control ♀ v mutant ♀ *; control ♀ v control ♂ *; control ♀ v mutant ♂ n.s.; mutant ♀ v control ♂ |
| | | | males control=5; mutant=3 | ~P30 | 6.9222 ± 0.314 | 5.089 ± 0.173 | | | |

| | | | | | | | | | |
|---|---|---|---|---|---|---|---|---|---|
| | | | | | | | | P = 0.6582; control ♂ v mutant ♂ P = 0.0050 | ****; mutant ♀ v mutant ♂ n.s.; control ♂ v mutant ♂ ** ; |
| | | | | | **Line of best fit male** | **Line of best fit female** | | | |
| Suppl. data 6c | Control weight kinetics slope | control males vs control females | | | y=1.658x +13.74 | y=1.270x +11.96 | Simple linear regression of slopes | P =0.0030 | ** |
| Suppl. data 6d | Mutant weight kinetics slope | mutant males vs mutant females | | | y=2.669x +4.811 | y=2.437x +6.407 | Simple linear regression of slopes | P =0.3404 | n.s. |
| Suppl. data 7a | GH (ng/ml) | control vs mutant | 6 | 13 weeks | 2.8517 ± 0.4573 | 0.7417 ± 0.1393 | Student's two-tailed T-test | P =0.0013 | * |
| | ACTH (pg/ml) | control vs mutant | 6 | 13 weeks | 35.45 ± 3.1135 | 52.5833 ± 7.7925 | Student's two-tailed T-test | P =0.0684 | n.s. |
| | PRL (ng/ml) | control vs mutant | 6 | 13 weeks | 59.9167 ± 17.1654 | 102.1667 ± 19.2153 | Student's two-tailed T-test | P =0.1321 | n.s. |
| | TRH (pg/ml) | control vs mutant | 6 | 13 weeks | 129.1667 ± 15.8017 | 115.1667 ± 15.7129 | Student's two-tailed T-test | P =0.5439 | n.s. |
| | TSH (ng/ml) | control vs mutant | 6 | 13 weeks | 0.47 ± 0.0786 | 2.13 ± 0.4565 | Student's two-tailed T-test | P =0.0050 | ** |
| | T4 (ng/ml) | control vs mutant | 6 | 13 weeks | 28.2333 ± 3.0032 | 16.7833 ± 1.0422 | Student's two-tailed T-test | P =0.0048 | ** |
